# Germline-somatic residue synergy reshapes antibody encounter-state pathways to enhance HIV-1 recognition

**DOI:** 10.64898/2026.03.26.714541

**Authors:** Sangita Kachhap, Yishak Bililign, Jared Lindenberger, Carrie Saunders, Priyamvada Acharya, Rory Henderson

**Affiliations:** Duke Human Vaccine Institute, Duke University Medical Center, Durham, NC 27710, USA; Department of Medicine, Duke University Medical Center, Durham, NC 27710, USA; Department of Surgery, Duke University Medical Center, Durham, NC 27710, USA

## Abstract

Antibodies initiate antigen recognition through short-lived encounter states. These states are structurally distinct from the bound state and allow an antibody to reach the epitope from diverse approach angles. We previously showed that encounter states enable broadly neutralizing HIV-1 antibodies to access protected epitopes, but how these states evolve during affinity maturation is unknown. Here, we defined encounter-state ensembles for two intermediates in a glycan-dependent antibody clonal lineage using extensive molecular dynamics simulations and Markov state modeling. Somatic mutations in the more mature member did not stabilize the bound state. Instead, they created early glycan-mediated interactions that reoriented the antibody during approach and markedly increased the association rate. This reorientation redistributed productive encounter states across a larger antigen surface. This expanded the antigen surface area over which collisions led to productive binding. The modified encounter state landscape positioned germline residues for conserved contacts and provided a kinetic route that naturally bypasses steric barriers at the epitope. Together, these results show that affinity maturation can proceed by reshaping encounter pathways rather than altering the final complex, revealing a generalizable mechanism by which somatic–germline synergy enhances antigen recognition.

## Introduction

Antibody-antigen affinity is related to interaction kinetics through the ratio of the dissociation and association rate constants. The dissociation rate constant provides a mechanistic context for determining factors that affect interaction lifetime, while the changes in the association rate constant can reveal limitations to recognition (1). Dissociation often governs specific residue-level interactions revealed in high-resolution structures. In contrast, association bottlenecks arise from steric and geometric constraints that slow the initial encounter, such as epitope-paratope shape mismatch, proximal loops or glycans, or conformational specificity (2, 3). Bottlenecks to association include epitope-paratope shape mismatch and steric hindrance from proximal loops, glycans, or antigen conformation specificity (4, 5). Comparing bound and unbound structures can identify many of these bottlenecks. However, as transition states are extremely transient, it is difficult to determine how antibodies overcome these barriers.

Bottlenecks to association are particularly acute for antibodies targeting the HIV-1 Envelope (Env) fusion glycoprotein due to its dense glycan shield and five highly sequence variable loops that protect conserved, surface-exposed motifs (6–11). Antibodies that overcome these obstacles to recognition and neutralize diverse HIV-1 viruses have been isolated from people living with HIV-1 (12–16). Referred to as broadly neutralizing antibodies (bnAbs), these antibodies are characterized by extensive somatic mutation, which enables broad recognition despite variation in epitope exposure. HIV-1 bnAbs target specific sites of vulnerability on the Env, including the host receptor CD4 binding site, the timer apex, and a co-receptor binding site located at the base of variable loop 3 (V3) (14, 17–19). Each site presents conserved Env residues that are protected sterically by proximal glycans and variable loops, which differ in sequence composition, length, and conformation (10, 20–24). Broadly neutralizing antibodies share common features, including long HCDR3 Loops, accumulation of rare somatic mutations, and shared gene usage (25–28). The ontogenies of these antibodies are among the most well-studied, with a wealth of liganded and unliganded epitope structures available to interrogate specific recognition bottlenecks.

The co-receptor binding site is vulnerable to bnAbs that form interactions with a conserved GDIR/K motif in the V3 loop and glycan at position 332, which is ∼74% conserved at this position (29, 30). These V3-glycan epitope-targeting bnAbs are characterized by long HCDR3 loops that penetrate the glycan shield, forming the major protein-protein contact surface. Heavy-chain gene usage is less restrictive than that of bnAbs targeting other epitopes, such as the CD4-binding site, resulting in greater variance in binding approach angles. Several bnAb families target this region, including lineages such as DH270 (31) and PGT128 (32), each with a distinct germline gene but showing similar reliance on long HCDR3 loops. Unlike many bnAbs targeting other epitopes, where glycans hinder antibody binding, the V3-glycan bnAbs make direct use of the glycan shield to facilitate binding (28, 33). Nevertheless, the glycan shield presents a substantial association bottleneck, especially at the germline and early maturation stages (34). Several immunogens designed to elicit these bnAbs by vaccination eliminate glycans in and near the epitope proximal V1 loop, with the V1 loop itself also being a barrier to association (34–36). The V1-loop sequence and sequence length vary markedly between isolates, often resulting in severe steric hindrance for V3-glycan bnAbs (4, 30, 37). Structures for these bnAbs show that breadth is a result of both direct glycan interactions and shifting of the V1 loop to accommodate antibody engagement of the GDIR/K motif.

The DH270 clonal lineage is among the most well-studied, with high-resolution bound state structures determined across the maturation pathway (4). The broadest and most potent clone antibody, DH270.6, neutralizes roughly 50% of viruses from a representative global virus panel (31). The clonal lineage is represented by two distinct clades that separated early during maturation after the first intermediate, referred to as I5.6. The clade containing DH270.6 proceeds from I5.6 through two intermediate clade members: the post-I5.6 intermediate I3.6, followed by intermediate I2.6 (4). Heterologous neutralization breadth first appeared in the I3.6 intermediate, marking a key stage in maturation. Of the twelve mutations that occurred between I5.6 and I3.6, two are critical for the acquisition of breadth in I3.6 (4): heavy chain variable domain (V_H_) R98T and light chain variable domain (V_L_) L48Y. Both mutations increase the number of contacts with the key epitope glycan at position 332 (4). These additional bound-state contacts did not lead to an enhanced dissociation rate for I3.6 relative to I5.6. Rather, the association rate constant improved by nearly 12-fold in I3.6 (31). This observation indicated that these mutations primarily impacted structural states involved in that association transition barrier. An examination of these structural states suggested that the V_H_ R98T and V_L_ L48Y mutations expanded the productive collision surface area on the Env (38). This was accomplished through rapid post-collision N332-glycan capture. This strong glycan interaction formed a tether that allowed the antibody to rotate around the glycan to reach the epitope. This binding mechanism explained the apparent disconnect between structure and kinetics measurements and led to improved vaccine immunogen engineering (38). However, these structures did not reveal how somatic mutations introduced during affinity maturation influenced these transition structures.

Here, we asked whether the intermediate antibodies I5.6 and I3.6 form encounter states similar to those of the mature DH270.6 bnAb, and how somatic mutations reshape the association pathway. We performed extensive adaptive-sampling molecular dynamics simulations and built Markov State Models (MSMs) to define the association pathways between each intermediate and an HIV-1 Env. Based on differences in their mutational profiles, we hypothesized that glycan capture and tethering would be inefficient in I5.6, thereby reducing the productive collision surface available to this early intermediate. Our results confirm that I5.6 fails to capture the N332 glycan early in the association transition, leading to encounter orientations that disfavor engagement of conserved epitope residues by germline-encoded HCDR3 motifs. In contrast, double-mutant cycle analysis showed that germline-encoded residues in I3.6 cooperate with somatic mutations to stabilize both association and dissociation transition states. Together, these results demonstrate that germline-encoded motifs and somatic mutations acquired during affinity maturation act synergistically in the DH270 lineage to enhance Env recognition.

## Results

### Molecular dynamics-based Markov state models identify I5.6 and I3.6 association paths to the bound state

We first asked whether the early and late intermediates in each antibody differ in how they reach the bound state, rather than how stable that state is. The DH270.6 association pathway involved the rapid capture and tethering of the N332-glycan D1 arm by V_H_ D115 and V_L_ Y48 following initial collisions near the V4 Loop (38). Tethering effectively expands the viable Env collision surface available to DH270.6, thereby enhancing association rates and reducing the impact of site-specific steric occlusion in highly sequence variable regions, such as V1 (Figure 1) (38). It is unclear whether this is a germline-encoded feature or acquired through somatic hypermutation. Since I3.6 acquired the V_H_ R98T, which frees D115 to form additional N332-glycan contacts and N332-glycan interactive V_L_ L48Y mutations, we reasoned that I3.6 N332-glycan capture and tethering efficiency would be higher than that of the earlier I5.6 intermediate, and is thus acquired during maturation. To assess these, we first mapped the I3.6 and I5.6 association pathways by simulating recognition of a CH848 HIV-1 Env known to form high-affinity interactions with both intermediates (4). Encounter state lifetimes are short, on the order of microseconds, and their structures are dynamic. These features render their study by traditional structural biology methods intractable. We and others have shown that extensive molecular dynamics simulations, paired with Markov State Models (MSMs), effectively describe encounter-state structure and exchange (38–42). To obtain transition structures between states, many cycles of Adaptive MD simulations were run, followed by MSM construction from successive MD trajectories using structural features that define the binding process. As structural features, we used distances between bound-state contacts observed in high-resolution structures. These features were used to define a kinetic structure mapping. The kinetic mapping was then used to identify a minimal subset of kinetically distinct structures/states and how they exchange (43, 44). The resulting MSMs show where the antibodies are positioned relative to the Env and which fast-exchanging structures cluster together. Standard modeling criteria for convergence were met for each antibody MSM, and the known bound-state conformation was correctly identified as the dominant population (Figure 1 and Supplementary Figures 1-5) (4). Recovery of the correct bound state demonstrates that the MSMs reliably capture the binding landscape.

**Figure 1.**
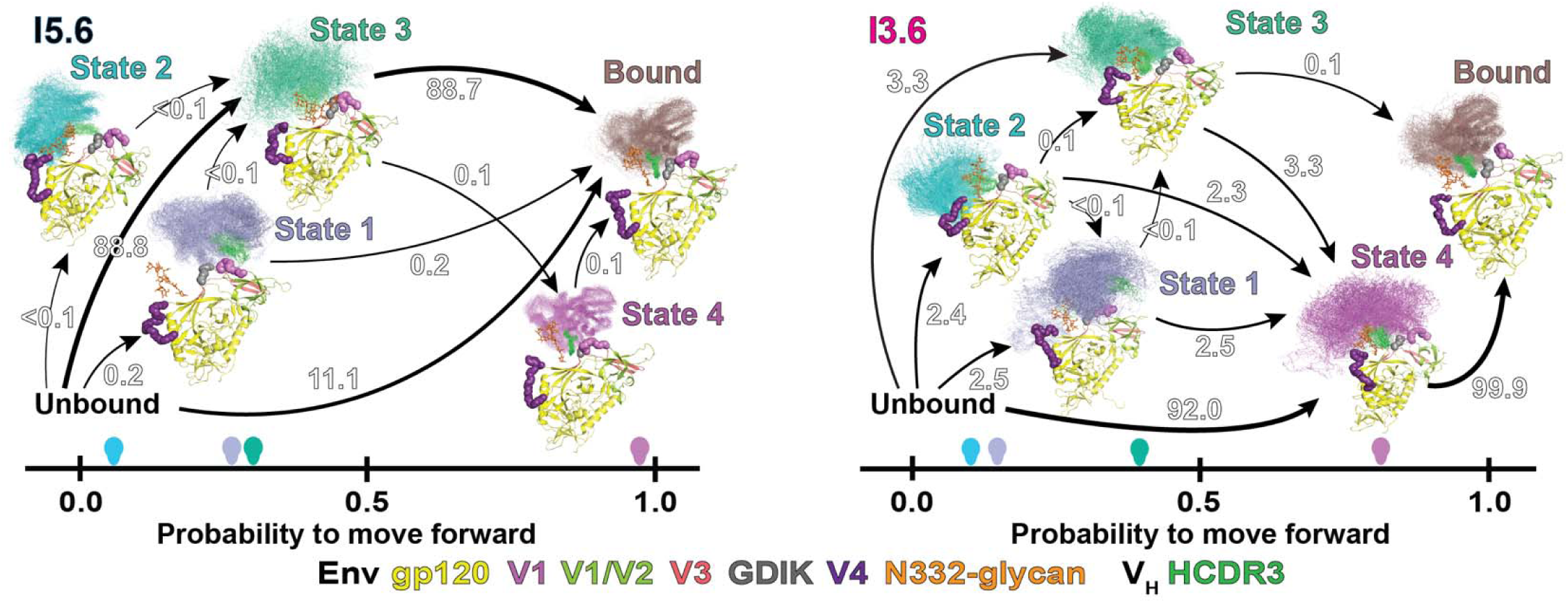
Association pathways for and I5.6 (*left)* and I3.6 (*right*) are represented by the net flux from the unbound to the bound state. Pathways are plotted along the forward committer probability, representing the probability to move forward from a given state. A total of 50 antibody V_H_/V_L_ domain structures are shown for each state, representing the state encounter-ensemble. Structures were sampled from the MSMs, weighted by their stationary state probabilities. Arrow colors indicate the committer probability for the corresponding state, based on the V_H_/V_L_ color of that state. Arrows indicate state connectivity for transitions toward the bound state with arrow weight representing the relative flux along transition paths. Numbers above each arrow indicate the percent contribution to the total transition flux for each transition.

### Increased encounter state epitope contact enhances binding probability

We next examined how each antibody reaches the bound state. The MSMs describe the time-dependent exchange between all simulated states. These include unbound, encounter, and bound states. We are interested in which encounter states form first, and how those states move toward the bound state. We used transition path theory (45) to define the MSMs in terms of the probability each encounter state transitions to the bound state and the number of transitions between states per unit time. We refer to this encounter mapping as the association pathway. The I5.6 and I3.6 intermediate association paths both displayed 4 distinct encounter states (Figure 1A). These include a V1-loop interactive state, State 1_V1_, a V4-loop proximal state, State 2_V4_, and an epitope GDIK motif proximal state, State 3 (Figure 1A). The I5.6 association path included a stable near-bound state, State 4_I5_, whereas the I3.6 association path included a dynamic prebound state, State 4_I3_ (Figure 1A). The unbound state had access to all encounter states in each pathway. Unlike I5.6, I3.6 did not reach the bound state directly. The I3.6 association pathway was characterized by enhanced utilization of encounter states and greater exchange among them. Conversely, successful I5.6 encounters were dominated by encounters formed near the bound state (Figure 1A). These results demonstrate somatic mutations in I3.6 are responsible for expanding the productive collision surface in the DH270 bnAb clone.

Across both antibody pathways, encounter position alone was insufficient to predict binding probability; instead, orientation-dependent epitope contact determined progression to the bound state. The encounter structure ensembles in each pathway differed in their overall antibody orientation relative to the epitope. Visual inspection of the State 1_V1_ encounters with the V1 loop in each antibody revealed marked differences in the HCDR3 loop orientation relative to the epitope (Figure 1A, Supplemental Figures 4 and 5). The HCDR3 loop is positioned near the V1 loop in the I5.6 association path with LCDRs 1 and 3 forming dynamic contact with the GDIK (Figure 1A, Supplemental Figure 4). In contrast, the I3.6 State 1_V1_ encounter does not form substantial GDIK motif contact, and the HCDR3 is oriented away from the V1 loop (Figure 1A, Supplemental Figure 5). The greater epitope residue contact by I5.6 State 1_V1_ is consistent with the greater probability that it will reach the bound state compared to I3.6 State 1_V1_ (Figure 1A). Like the V1-loop encounter state, while both I5.6 and I3.6 form a V4-loop proximal state, State 2_V4_, the HCDR3 orientation relative to the epitope differs substantially, with I3.6 positioning the HCDR3 loop toward the GDIK motif and I5.6 orienting its HCDR3 loop away from the epitope (Figure 1A, Supplemental Figures 4 and 5). Greater epitope contact in I3.6 in State 2_V4_ is, like greater epitope contact in I5.6 State 1_V1_, also associated with a higher probability of transitioning to the bound state. The State 3 encounter positions the antibodies in closer proximity to the GDIK epitope motif. However, I5.6 exhibits greater dynamics in this state and positions the antibody farther from the epitope, resulting in a reduced probability of transitioning to the bound state (Figure 1A and B, Supplemental Figures 4 and 5). In each encounter state, greater encounter state epitope contact enhances the probability that the encounter proceeds to the bound state.

### Somatic mutations in I3.6 expand the productive Env encounter surface

In principle, I5.6 and I3.6 have equal access to all encounter surfaces on the Env. The primary differentiator is thus the degree to which each antibody utilizes a given encounter state at a particular Env position. The frequency of transitions from encounter to bound states can be used to measure utilization. We asked whether somatic mutations in I3.6 increased the number of productive binding transition events through each encounter state. The percent encounter state utilization was defined here as the number of crossings from a given state to the next per 100 ns, referred to as the net reactive flux, relative to the total reaction flux. This yields the reactive flux defined as the percent contribution of each encounter transition to the full association pathway. The I5.6 association path primarily flowed through epitope GDIK motif proximal State 3, accounting for 88.7% of the reactive flux, or through direct formation of the bound state, accounting for 11.1% of the reactive flux (Figure 1A, Supplemental Figure 1). The V1-contact and V4-proximal encounter state formation represented <0.1% and 0.2% of the reactive flux, respectively, and reach the bound state through State 3, which predominantly transitions directly to the bound state. A minority of the flux to the bound state occurs through the prebound State 4_I5.6_ intermediate (Figure 1A, Supplemental Figure 1). These results show that the I5.6 association is restricted to a narrow collision surface near the epitope. Thus, although I5.6 is capable of glycan engagement in the bound state, it fails to deploy this interaction early enough to influence encounter-state flux.

Despite coarse structural similarity between the I5.6 and I3.6 association pathway states, the reactive pathways differ markedly. The I3.6 association path distributes reactive flux more evenly across the intermediate encounter states without directly accessing the bound state, as occurred in the I5.6 association path (Figure 1A, Supplemental Figure 2). The dominant transition passes through the pre-bound State 4_I3.6_ before reaching the bound state, accounting for 91.9% of the reactive flux. Flux through States 1_V1_ and 2_V4_ is enhanced relative to I5.6, at 2.5% and 2.4% each, with State 2 _V4_ gaining access to States 1_V1_ and 4_I3.6_ in addition to flux through State 3. Most flux through State 2 _V4_ transitions to the bound state through State 4 _I3.6_. Similarly, State 1_V1_ gains access to the prebound State 4 _I3.6_, while retaining flux through State 3, similarly focusing flux through State 4 _I3.6_. States 3 and 4 _I3.6_ both access the bound state directly, with passage through State 4 _I3.6_ accounting for most of the reactive flux, at 99.9% (Figure 1A, Supplemental Figure 2). These results show that the I3.6 somatic mutation-induced changes in encounter orientation alter encounter-state utilization, enabling more effective use of epitope-distant states. This creates a larger Env surface target for productive I3.6 collisions, explaining the enhanced I3.6 association rate relative to I5.6 (38).

### Somatic mutations in I3.6 enable epitope-glycan capture in encounter states

The mature DH270.6 bnAb forms extensive N332-glycan contact in encounter states through V_H_ D115 and V_L_ Y48 contact with the glycan D1-arm. High-resolution bound state structures show that the I5.6 intermediate forms glycan D1-arm interactions as well. This suggests I5.6 has the capacity to form these interactions in the encounter states. However, an immunogen designed to enhance DH270.6 encounter-state exposure and transition to the bound state did not appreciably increase I5.6 association rates, despite substantial rate improvements in I3.6. Given the minimal reactive flux through distant encounter states in I5.6, we hypothesized that I5.6 fails to form N332-glycan interactions in encounter states. The bound state DH270 N332-glycan interactions are primarily formed between residues in or near a cleft formed at the V_H_/V_L_ interface at the HCDR3 loop base (Supplemental Figure 6A) (4). To determine the extent of N332-glycan capture and tethering in each encounter state, we determined the number of contacts formed between the N332-glycan D-arm Mannose residues and antibody contacts observed in published I5.6 and I3.6 structures (Supplemental Figure 6A). The State 1_V1_ encounters showed relatively few contacts per antibody, as expected given the antibody orientations in each state (Supplemental Figure 6B). The I3.6 State 2_V4_ displayed a substantially larger contact count relative to I5.6 (Figure 2A). This is consistent with inspection of the encounter state structure ensembles from the I3.6 and I5.6 State 2_V4_, which show that, despite proximity, the N332-glycan is oriented away from I5.6 while it is oriented toward I3.6 (Supplemental Figure 6C). The number of N332-glycan contacts increased in State 3 in both I3.6 and I5.6, with the biggest gains in I5.6 (Figure 2B), which nearly reaches parity with I3.6. The contacts are nevertheless fewer and more variable in the I5.6 State 3 (Figure 2B). The number of contacts increases further for both antibodies in their respective State 4 states, with I5.6 displaying a larger number of contacts (Figure 2C). This difference is consistent with the distinct encounters formed in each association path, with State 4_I5.6_ occupying an almost completely bound state and State 4_I3.6_ occupying a pre-bound transition intermediate (Figure 1A). These results show that the I3.6 somatic mutations enable encounter-state glycan capture, thereby more effectively tethering the glycan during the transition to the bound state.

**Figure 2.**
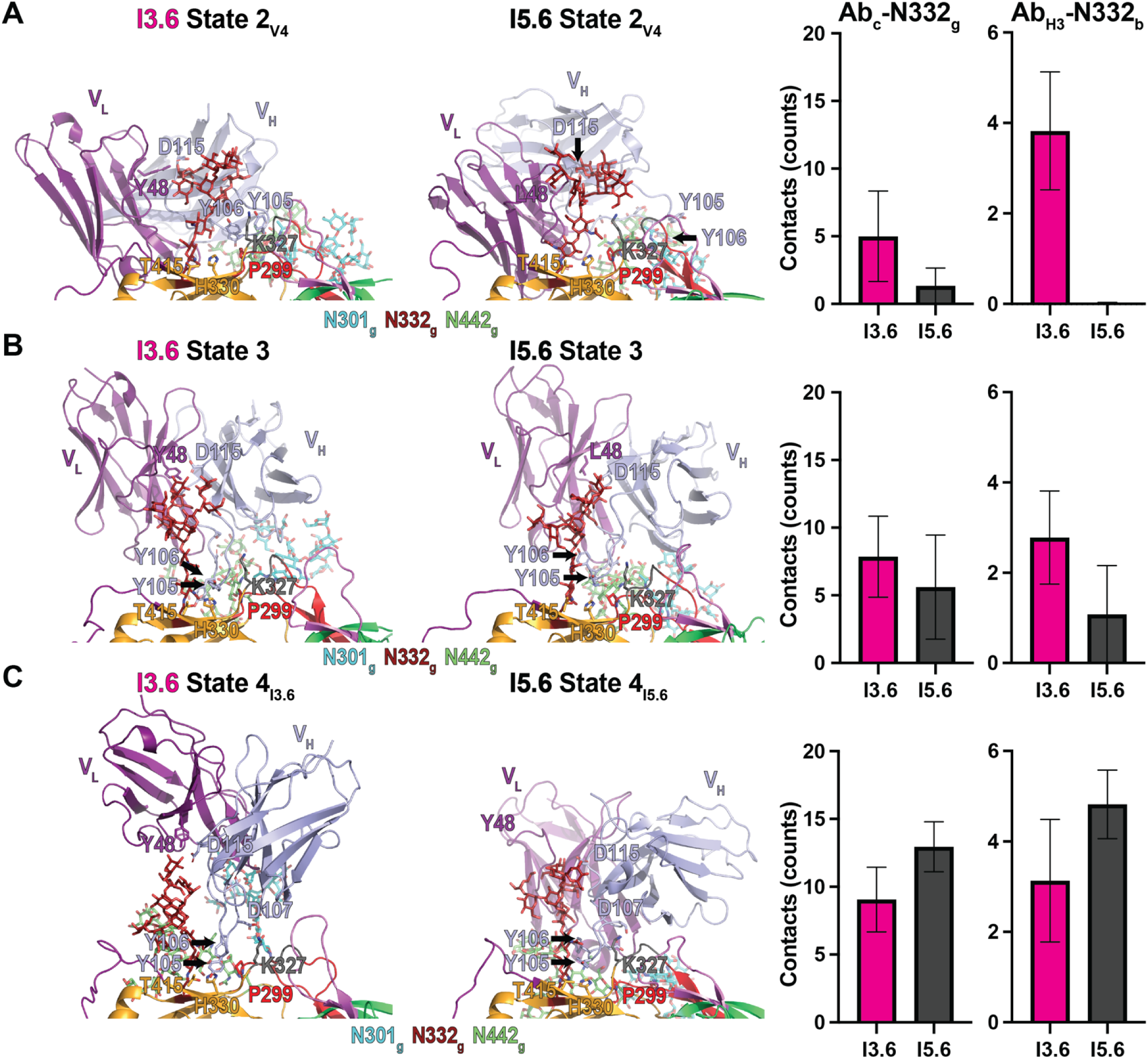
**(A)** (*left*) Representative State 2 structure identified in the I3.6 and I5.6 MSMs. (*right*) Contact counts between I3.6 and I5.6 V_H_/V_L_ antibody cleft and cleft proximal residues (Ab_c_) with N332-glycan (N332_g_) D-arm residues and between antibody HCDR3 (Ab_H3_) residues and residues near the N332-glycan base (N332_b_). **(B)** (*left*) Representative State 3 structure identified in the I3.6 and I5.6 MSMs. (*right*) Contact counts between I3.6 and I5.6 V_H_/V_L_ antibody cleft and cleft proximal residues (Ab_c_) with N332-glycan (N332_g_) D-arm residues and between antibody HCDR3 (Ab_H3_) residues and residues near the N332-glycan base (N332_b_). **(C)** (*left*) Representative State 4 structure identified in the I3.6 and I5.6 MSMs. (*right*) Contact counts between I3.6 and I5.6 V_H_/V_L_ antibody cleft and cleft proximal residues (Ab_c_) with N332-glycan (N332_g_) D-arm residues and between antibody HCDR3 (Ab_H3_) residues and residues near the N332-glycan base (N332_b_). Contacts represent the state MSM weighted mean number of contacts, defined as residues within 4.0 Å. Error bars represent the state MSM weighted contact standard deviation.

### Epitope-glycan capture enables early bound state contact formation in the I3.6 association path

The DH270 HCDR3 is generally permissive to substitutions (28). However, key paratope residues are essential for germline recognition of the Env. These include an HCDR3 ^105^YYD^107^ motif encoded by the antibody germline gene and a proximal germline S109_Ab_ residue (4, 28). These residues make extensive contact with the glycan and protein portions of the Env epitope (4, 35). We therefore asked whether contact between the ^105^YYD^107^ and S109_Ab_ residues with the epitope influenced the encounter transitions (Figure 1A). The change in antibody orientation observed in the I3.6 State 2_V4_ encounter oriented the HCDR3 toward the DH270 epitope. Examination of the I3.6 State 2_V4_ encounter state structures revealed dynamic contacts formed between the ^105^YYD^107^ motif and several epitope or epitope-proximal Env residues. These included residues P299_Env_, K327_Env_, H330_Env_, T415_Env_, and Q417_Env_ (Figure 2A, Supplemental Figure 6D). The I3.6 contacts are primarily mediated by interactions between HCDR3 Y105_Ab_ and gp120 P299_Env_ and K327_Env_, as well as by HCDR3 Y106_Ab_ interaction with Env residue P299_Env_ (Supplemental Figure 7A and B). This positions the HCDR3 loop at the N332-glycan base, such that antibody rotation around the glycan brings the antibody into contact with the epitope.

Unlike I3.6, the I5.6 State 2_V4_ does not form HCDR3 contact with the Env protein surface. Transition to the bound-state proximal State 3 in I5.6 yields a diffuse encounter-state ensemble that lacks the specific surface anchoring observed in I3.6 (Figure 1). Transitions to State 3 reduced the average number of HCDR3 contacts in I3.6 relative to State 2_V4_ and represent the first appearance of epitope and epitope-proximal contacts in an I5.6 encounter (Figure 2A and B). State 3 is the dominant encounter intermediate in the I5.6 association path. In I3.6, the State 3 orientation is comparatively fixed and accounts for a minority of the reactive flux. Structural overlap with the State 4_I3.6_ encounter suggests the I3.6 State 3 is a long-lived state along the rotation path to the epitope. The loss in contacts in I3.6 in State 3 relative to State 2_V4_ is related to the elimination of non-native bound state contacts in the State 2_V4_ encounter. Losses include I3.6 HCDR3 Y105_Ab_ and gp120 K327_Env_ and Y106_Ab_ contact with P299_Env_ in State 3 (Supplemental Figure 7A and B). Native-like contact with H330_Env_ is gained in the I3.6 State 3 (Supplemental Figure 7A). These contact changes show that I3.6 begins forming native contacts early in the binding event. Conversely, the I5.6 intermediate gains limited contact between HCDR3 Y105_Ab_ and gp120 P299_Env_, K327_Env_, and H330_Env_ (Figure 2B and C, Supplemental Figure 7A). Contact between I5.6 HCDR3 Y105_Ab_ and gp120 H330_Env_ is similarly limited in State 3 (Supplemental Figure 7A). This results in less native-like encounter contact relative to the I3.6 State 3. These results show that the HCDR3 orientation shift in the I3.6 encounters enables early formation of native-like contacts.

The final encounter states in each intermediate association path differ substantially. State 4_I5.6_ is a minor encounter state that is nominally bound, forming stable contacts between HCDR3 Y105_Ab_ and gp120 H330_Env_, T415_Env_, and Q417_Env_ in addition to Y106_Ab_ and D107_Ab_ contact with gp120 K327_Env_ (Figure 2C, Supplemental Figure 6D and 7A, B, and C). The State 3_I3.6_ is the dominant unbound-to-bound encounter intermediate in the I3.6 association path. The state exhibits wide structural variability while maintaining a bound-state contact profile. Contacts include HCDR3 Y105_Ab_ interaction with gp120 H330_Env_, along with T415_Env_ and Q417_Env_ interactions and elimination of non-bound state contact with P299 (Figure 2C, Supplemental Figure 6D and 7A). State 4_i3.6_ also acquires HCDR3 D107_Ab_ contact with gp120 K327_Env_ (Supplemental Figure 7C). Binding was preceded by this non-native contact in the DH270.6 association path (38). These results show that somatic mutations in I3.6 that enabled encounter state recognition of the N332-glycan also enabled germline-encoded recognition of key bound-state contacts early in the transition.

### Critical germline encoded HCDR3 residues play dual association and dissociation roles in I3.6

The association pathway in I3.6 shows that encounter states acquire native-bound-state contacts. These include contacts between key HCDR3 residues that enable germline Env recognition. This suggests these residues play important roles in both association and dissociation. Despite contact between these residues in encounter states, the precise three-dimensional arrangements are not yet formed. In the bound state, HCDR3 residues Y105 and Y106 flank the two N332-glycan GlcNAc residues, packing against the first and second GlcNAc residues, respectively, with the potential to form CH−π interactions (Supplemental Figure 8) (4). Two additional key HCDR3 residues, D107_Ab_ and S109_Ab_, form a networked interaction with the conserved Env GDIK/R motif through the V_H_ Y33 residue and a structural water molecule (Supplemental Figure 8). This complex interaction network is not yet formed in the encounter states. Thus, the association and dissociation processes are likely mechanistically distinct. We previously showed that I3.6 gains over an order-of-magnitude improvement in association rate constant (k_a_) relative to I5.6, at 1.9×10^3^ and 1.6×10^2^ M^-1^s^-1^, respectively, for binding to an autologous Env that binds I5.6, referred to as CH848.d949 (4). Two distinct modifications were made to this Env to enhance the association rate for I3.6 (38). The first eliminates a PNGS in the V4 loop at position 407, opening the State 2_V4_ encounter surface to collision. The second eliminates a PNGS position 442. Both resulted in a marked enhancement in the association rate in I3.6 and I5.6. The I3.6 Fab dissociation rate constant was, however, unaffected by these substitutions (38). These results are consistent with the association paths shown here, indicating that I5.6 and I3.6 have access to the State 2_V4_ encounter state, and that substitutions that facilitate association transitions do not necessarily affect dissociation rates. The results also indicate that the key HCDR3 residues play an essential role in determining the impact of encounter states on association and in anchoring the antibody to the epitope in the bound state (4, 38). We therefore asked whether the HCDR3 residues play a dual role in I3.6 binding, enhancing association through encounter contacts as predicted by the association paths, and maintaining the bound state through interactions with the GDIK motif and N332 glycan.

To compare HCDR3 residue impacts on association and dissociation rates in the I3.6 intermediate, we measured binding kinetics using surface plasmon resonance (SPR) for alanine substitutions at HCDR3 residues Y105 _Ab_, Y106 _Ab_, D107 _Ab_, and S109_Ab_ (Table 1, Figure 3A, Supplemental Figure 9). The mean association rate constants (k_a_) were 2.4×10^3^, 8.5×10^2^, 8.1×10^2^, 9.4×10^2^, and 1.2×10^3^ M^-1^s^-1^ for the unmutated I3.6 Fab and the Y105A_Ab_, Y106A_Ab_, D107A_Ab_, and S109A_Ab_ mutants, respectively. Approximately 2-3 fold reductions in the association rates were observed for the mutants relative to the unmutated Fab (Table 1, Figure 3A, Supplemental Figure 9A). The mature DH270.6 Fab k_a_ was similar to the I3.6 Fab at 1.7×10^3^ M^-1^s^-1^. Changes in the dissociation rate constants (k_d_) were substantial with the I3.6 k_d_ lower than the calibrated instrument detection limit of 7.0×10^-6^ s^-1^ and the Y105A _Ab_, Y106A _Ab_, D107A _Ab_, and S109A_Ab_ at 4.2×10^-3^, 1.0×10^-4^, 7.0×10^-4^, and 1.7×10^-4^ s^-1^, respectively, for fold changes of 600, 14, 100, and 24 relative to the detection limit, respectively (Table 1, Figure 3B, Supplemental Figure 9B and C). Like the I3.6, the DH270.6 k_d_ was lower than the detection limit of 7.0×10^-6^ s^-1^ (Table 1, Figure 3B, Supplemental Figure 9B and D). These results are consistent with encounter-state observations, indicating that these residues play a role in association and dissociation, and show that the mature antibody, though substantially more broadly neutralizing, does not exhibit rate constant enhancement for CH848.d949 SOSIP Env.

**Figure 3.**
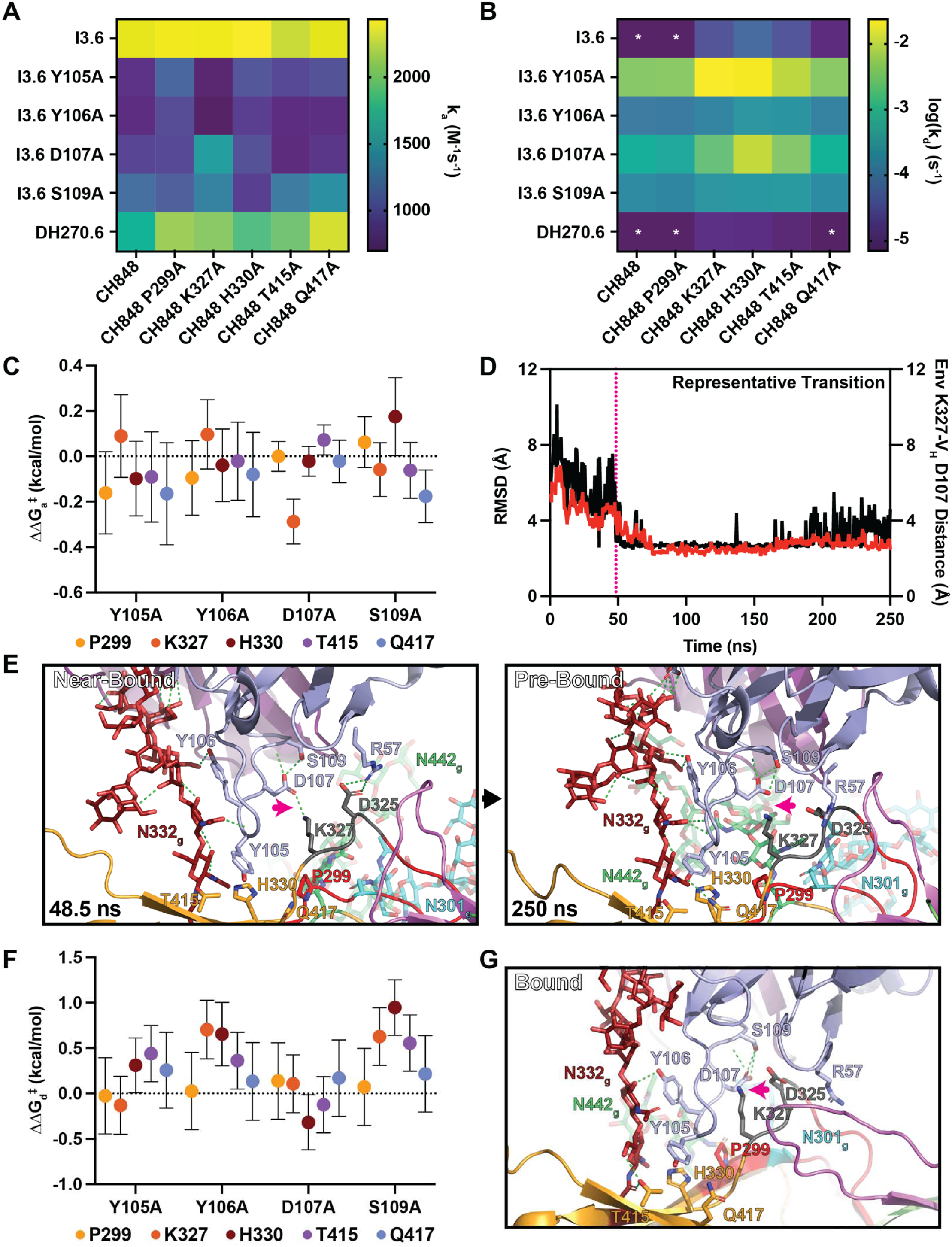
**(A)** Mean association rate constant value (n=3) heatmap for pairwise interactions between the I3.6 intermediate, I3.6 alanine mutants, and the mature DH270.6 antibody Fabs and CH848.d949 SOSIP and CH848.d949 alanine mutant SOSIPs. **(B)** Log mean dissociation rate constant value (n=3) heatmap for pairwise interactions between the I3.6 intermediate, I3.6 alanine mutants, and the mature DH270.6 antibody Fabs and CH848.d949 SOSIP and CH848.d949 alanine mutant SOSIPs. Asterisks indicate dissociation rate constants below the calibrated instrument detection threshold (<7e-6 s^-1^). Colors for these rate constants were assigned based on the threshold value. **(C)** Double mutant cycle free energy differences in the association transition barrier free energy between pairwise I3.6 HCDR3 and CH848.d949 epitope residues. Error bars indicate the standard error of the mean. Errors were propagated from sensorgram fit and technical replicates (n=3) for each measurement set used in calculations. **(D)** Representative simulation depicting the transition from a near-bound to a prebound state. The root mean square deviation (RMSD; red) is plotted on the left y-axis. The distance between HCDR3 residue D107 γ-carbon and K327 ϵ-amine is plotted on the right y-axis. The pink dashed line indicates the 48.5 ns timepoint at which the stable D107-K327 interaction forms. **(E)** (*left*) Near-bound encounter state structure from the representative transition in (*D*) at the 48.5 ns timepoint. The pink arrow identified the D107 interaction with K327. (*right*) Pre-bound encounter state structure from the final trajectory state at the 250 ns timepoint. The pink arrow identifies the absence of an interaction between D107 and K327. **(F)** Double mutant cycle free energy differences in the dissociation transition barrier free energy between pairwise I3.6 HCDR3 and CH848.d949 epitope residues. Error bars indicate the standard error of the mean. Errors were propagated from sensorgram fit and technical replicates (n=3) for each measurement set used in calculations. **(G)** The DH270 bound state structure (PDB ID 8SB1). The pink arrow identifies the absence of an interaction between D107 and K327.

To examine impacts on kinetics related to epitope and epitope proximal residue substitutions, we measured the binding of I3.6 Fab and each I3.6 alanine mutant to the CH848.d949 SOSIP trimer with and without individual alanine substitutions at residues P299_Env_, K327_Env_, H330_Env_, T415_Env_, and Q417_Env_ (Table 1, Figure 3A, Supplemental Figure 9). These substitutions were measured for binding to the unmutated I3.6 Fab. Unlike the I3.6 Fab alanine substitutions, the SOSIP alanine substitutions did not show differences in k_a_ measured against the unmutated I3.6 Fab, with a mean k_a_ of 2.4×10^3^ ± 5×10^1^ M^-1^s^-1^ across the unmutated SOSIP and the alanine SOSIP mutants (Table 1, Figure 3A, Supplemental Figure 9A and C). The dissociation rate constants, however, showed substantial shifts for the K327A_Env_, H330A_Env_, T415A_Env_, and Q417A_Env_ substitutions, yielding 4.5, 8.6, 4.7, and 1.6 fold increases in k_d_ relative to the unmutated I3.6 vs. CH848.d949 detection limit of 7.0x10^-6^ s^-1^, respectively (Table 1, Figure 3B, Supplemental Figure 9). The P299A_Env_ mutant k_d_ was at the detection limit (Table 1, Figure 3A and B, Supplemental Figure 9B and D). These results show that the I3.6 intermediate relies heavily on the Y105_Ab_, Y106_Ab_, D107_Ab_, and S109_Ab_ HCDR3 residues to recognize and maintain an interaction with the CH848.d949 Env through key conserved Env residues.

### Alanine substitutions in I3.6 and CH848.d949 have minor impacts on fold stability

We next asked whether differences in kinetic rate may be due to changes in protein stability rather than in the binding mechanism. We measure thermal inflection temperatures (T_i_) via differential scanning fluorimetry (DSF) for each antibody and SOSIP alanine substitution. The I3.6 antibody Fab T_i_ was 78.8 °C with the Y105A_Ab_ mutation showing a ∼1 °C T_i_ increase and Y106A_Ab_ and S109A_Ab_ showing a ∼1 °C decrease (Supplemental Figure 10A). The D107A_Ab_ mutation showed a ∼3 °C reduction in T_i_ (Supplemental Figure 10A). The CH848 d949 SOSIP T_i_ values were unperturbed in the T415A_Env_ and Q417A_Env_ mutants relative to the unmutated form, each with Ti values of 73.6 °C (Supplemental Figure 10B). Reductions in T_i_ of less than 1 °C were observed for the P299A_Env_, K327A_Env_, and H330A_Env_ mutants (Supplemental Figure 10B). The association and dissociation rate constants did not display clear relationships with the antibody Fab or SOSIP T_i_ measurements (Supplemental Figure 11). The presence of discernible, though modest, differences in DSF thermal heating inflection temperatures indicates that the alanine substitutions impact the unfolding transition. We are, however, unable to establish a systematic assessment of the impact these mutations may have on local folded-state dynamics and on how such dynamics may affect bound-state stability.

### Long-range interactions affect association and dissociation transition-state free energies

We next asked whether germline-encoded HCDR3 residues that engage epitope elements during encounter states also govern bound-state stability, or whether these roles are mechanistically distinct. Direct interactions between paratope and epitope residues are often the primary focus of factors affecting affinity. However, long-range, non-contact residue interactions in encounter and bound states play essential roles in recognition and maintenance of the bound state. Although these interactions can be inferred from high-resolution structures or encounter state contacts, evaluating their impact on the binding free energy is challenging. Double mutant cycles (DMCs) provide a robust framework for quantifying the coupling free energy between interactive residues, regardless of distance (46–48). Residues are coupled when the change in binding free energy for both mutations measured together differs from the sum of the single-point mutants (49, 50). To measure residue coupling between key germline and epitope residues, we measured rate constants for the mutated antibody Fabs relative to the mutated SOSIP Envs. We are interested in the factors affecting association and dissociation processes separately. We therefore calculated transition barrier residue coupling using transition-state free energies derived from kinetic rate constants (49–52). Transition barrier coupling free energy differences using the following equation:

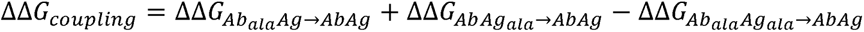

Each term represents the change in free energy going from the unmutated antibody-Env interaction to the alanine-substituted antibody, antigen, or both. A negative ΔΔG*_coupling_*value indicates that coupling lowers the transition barrier, while a positive value indicates that coupling raises the transition barrier. Because barrier heights determine kinetic rate constants, a lower transition barrier increases the transition rate in either direction. This increases association rates, thereby enhancing affinity. However, a lower dissociation transition barrier increases the dissociation rate, resulting in weaker overall binding affinity. Thus, affinity is enhanced by a negative coupling energy in the association phase and by a positive coupling energy in the dissociation phase. We use these measures to determine whether the I3.6 HCDR3 motif residues are coupled to the epitope-proximal residue patch during association. We then examine the role each residue plays in the dissociation barrier to determine whether these residues have mechanistically distinct effects on affinity.

### Germline HCDR3 residues lower the binding path activation barrier through an epitope patch

We considered a residue pair to be coupled when the estimated ΔΔG*_coupling_* differed from zero beyond the propagated measurement error. The I3.6 association pathway extensively used the HCDR3 ^105^YYD^107^ and S109_Ab_ motifs as anchors for glycan rotation. Such dynamic interactions are often present in encounter states, resulting in the spread of interactive energy across multiple residues. Experimental variance can, thus, mask pairwise residue coupling. The I3.6 HCDR3 motif interactions were distributed across several Env epitope and epitope-proximal residues in the encounter states, including Env residues P299_Env_, K327_Env_, H330_Env_, T415_Env_, and Q417_Env_ (Figure 2). We asked whether these interactions lower the transition free energy barriers for each I3.6 Fab alanine substitution binding to the unmutated Env. Association rate constants showed two to three-fold reductions relative to the unmutated Fab (Table 1, Figure 3A, Supplemental Figure 9A). The corresponding association barrier-free energies therefore increased from 0.4 to 0.6 kcal/mol for each ^105^YYD^107^ and S109_Ab_ residue substitution. We first quantified the impact the Env residue network had on each HCDR3 motif residue. The Y105_Ab_ and D107_Ab_ network couplings were -0.43 and -0.26 kcal/mol, accounting for 70.0 and 48.0 percent of the change in association barrier heights related to each alanine substitution, respectively. The Y106_Ab_ and S109_Ab_ residues did not exhibit substantial Env-encounter residue network couplings. These results show that observed encounter contacts between HCDR3 residues Y015_Ab_ and D107_Ab_ in I3.6 and Env play a key role in lowering the association transition barrier.

The Y105_Ab_ and D107_Ab_ HCDR3 residues showed coupling to the Env encounter interactive patch. The Y106_Ab_ and S109_Ab_ residues, however, did not. This indicates that the reduced association rates observed for these two residues are not distributed exclusively to the tested interactive patch residues. The residues may nevertheless show direct coupling with specific residues in the patch. We next evaluated the individual I3.6-to-Env residue ΔΔG*_coupling_* energies to determine whether specific residue pairs are coupled. Despite Y105_Ab_ coupling to the Env residues as a whole, no single Y105 _Ab_-Env residue pair displayed a ΔΔG*_coupling_* value greater than zero beyond the propagated measurement error. This suggests the coupling energy is distributed among the patch residues. The D107_Ab_ ΔΔG*_coupling_* energies were all near zero except for D107_Ab_-K327_Env_, which displayed a coupling energy of -0.3 kcal/mol. The K327_Env_ residue interacts with V_H_ Y33 and the backbone of HCDR3 residue S103 in the bound state (4). However, D107_Ab_ and Env K327_Env_ residues do not form direct contact in the bound state (4). We previously showed that an interaction between these two residues occurs during the transition from the intermediate to the bound state in the DH270.6 association pathway (38). This transition-related interaction was also observed in I3.6 simulations (Figure 3D). A representative transition from a near-bound MD trajectory shows D107_Ab_ interacting with K327_Env_ before the antibody rotates into a near-bound state. This state is eliminated as the antibody moves toward the fully bound state (Figure 3E).

The Y106 residue did not show patch coupling. This was reflected in the individual Y106_Ab_-Env residue pair that clustered near zero. This indicates the reduction in association rate in Y106_Ab_ is not attributable to coupling with the tested Env residues. The Y106_Ab_ residue showed encounter interactions only with P299_Env_ and only in State 2_V4_ (Supplemental Figure 7B). The N332-glycan is the primary Y106_Ab_ contact in both the encounter and bound states. The glycan interaction may therefore dominate the effects of the association barrier due to Y106_Ab_. The S109_Ab_ residue is the most distant epitope among the studied HCDR3 residues here. In the bound state, S109_Ab_ stabilizes the D107_Ab_ sidechain, which is positioned to form an extensive interaction network with adjacent antibody residues and a structural water. S109_Ab_ patch coupling was not observed beyond propagated error. However, coupling between I3.6 S109_Ab_ and Env H330_Env_ and Q417_Env_ was observed. Unlike the favorable coupling between Y105_Ab_ and D107_Ab_ and Env patch residues, the S109_Ab_ coupling with H330_Env_ was positive, indicating the interaction increases the association barrier. Conversely, Q417_Env_ coupling was negative, indicating that this coupling reduces the association barrier, thereby enhancing the on-rate. This long-range coupling suggests that S109_Ab_ influences HCDR3 structure and/or dynamics that impact HCDR3 interactions with H330_Env_ and Q417_Env_ during encounter states or encounter-state transitions. Together, the association rate couplings observed here show that a complex coupling network between the I3.6 HCDR3 residues and the Env epitope-proximal patch facilitates binding.

### Germline HCDR3 residues stabilize the bound state by mechanisms distinct from their binding path roles

Residue substitutions between interactive proteins can impact association, dissociation, or both. For residues that have dual impacts, their mechanistic roles in each process can be distinct. We next examined the transition-state coupling free energies for transitions out of the bound state, as defined by the dissociation rate constants. This allows us to compare coupling patterns that impact dissociation with those that impact association. Several constructs remained bound throughout the dissociation phase of our SPR measurements. This prevents accurate estimates of the dissociation rate constant. We used our calibrated detection limit, k_d_, of 7x10^-6^ s^-1^ for measurements at the limit in double-mutant cycles. The resulting ΔΔG*_coupling_* energies are therefore bounded estimates of the coupling strength. We observed bound state stabilizing coupling between I3.6 Y105_Ab_ and Env T415 _Env_, I3.6 Y106_Ab_ and Env K327_Env_, H330_Env_, and T415_Env_, and S109_Ab_ K327_Env_, H330_Env_, and T415_Env_ (Figure 3F). The mean coupling strengths varied from 0.5 to 1.0 kcal/mol. They were strongest for the I3.6 S109_Ab_ mutant, which may be related to the role this residue plays in stabilizing a structural water molecule that mediates HCDR3 backbone contact at an HCDR3 turn motif that contains the Env H330_Env_ and T415_Env_ interactive Y105_Ab_ residue and K327_Env_ contact. The I3.6 Y105_Ab_ and D107_Ab_ residues displayed coupling estimates greater than and less than zero within the propagated uncertainty, respectively. The remaining pairs did not show differences from zero within the propagated error. These results show that non-contact interactions between antibody and Env residues (Figure 3G) play an important role in both association and dissociation.

## Discussion

These results demonstrate that antibody affinity maturation can enhance antigen recognition by reshaping encounter-state pathways rather than only stabilizing the final complex. The results here show that somatic mutations in the DH270 clonal lineage expand the viable Env binding collision surface area. This occurs through direct, mutation-induced glycan tethering early in the association process, which in turn optimizes contact between germline residues and epitope residues throughout the association pathway. As in the DH270.6 association pathway (38), I3.6 forms N332-glycan D-arm contacts quickly after collisions spanning from the V1-loop to the V4-loop. Comparatively, the poor utilization of State 1_V1_ and State 2_V4_ in the I5.6 association path effectively reduces the viable collision surface to primarily epitope residues. This, coupled with low bound-state proximal State 3 net flux into the pre-bound a bound state compared to the I3.6 intermediate, explains its smaller CH848.d949 association rate constant. The tethering and induced HCDR3 contact with bound-state contact residues also likely explain the observed acquisition of heterologous breadth, beginning at the I3.6 intermediate (4). The N332-glycan is a critical neutralization sensitivity signature and is ∼74% conserved among HIV-1 viruses (53). The glycan itself is predominantly high-mannose and is, therefore, chemically conserved when present (54). Thus, tethering activity is likely retained regardless of variation in epitope sequence. The H330 and T415 residues are 83% and 80% conserved, respectively, and are also associated with neutralization sensitivity, while common substitutions at these sites are associated with resistance (55). With >98% P299 and GDIK/R motif K/R327 conservation, the encounter contacts used by neutralizing antibodies change little among neutralization-sensitive viruses. The association pathway is therefore likely conserved among HIV-1 Env isolates, as observed in bound-state structures.

The V1-loop poses a major hurdle for DH270.6 and its clone members (38). The effect is primarily steric occlusion of the epitope, a barrier that varies substantially depending on V1-loop length and glycosylation (4). The conserved nature of the I3.6 association path, shared by DH270.6 (38), and, in particular, the State 2_V4_-initiated path, is mechanistically favorable for limiting the effects of V1-occlusion on antibody association with Env. Unlike the State 1_V1_ encounter that forms direct V1-loop interactions, States 2 through 4 in I3.6 acquire nearly all bound state contacts before V1-loop engagement and displacement. The final step involves an HCDR3 D107 encounter with GDIK motif K327, which enables R57 to contact D325. This R57-D325 contact initiates the sweeping motion of R57 beneath the V1 Loop, thereby prying it off the epitope (4). Thus, while the V1-loop remains a barrier to recognition, the transition bottleneck it presents is minimized. Arginine residues similar to the DH270 R57 that contact at or near the GDIK/R motif are common among V3-glycan targeting bnAbs, and may represent a converged solution to V1-loop occlusion (35, 37, 56). For example, arginines at positions R94 and R100A in PGT121 and PGT128, respectively, form direct contact with the GDIK/R motif, displacing the V1 loop relative to its unliganded epitope position (37, 56). This suggests V1 loop displacement is an essential feature of this bnAb epitope class that can be overcome before reaching the fully bound conformation.

The I5.6 and I3.6 association pathways showed that somatic mutations and germline encoded residues play an essential, synergistic role in Env recognition and maintenance of the bound state. Our previous results demonstrated that removing barriers to State 2_V4_ formation and transition to the pre-bound state facilitated enhanced association through the I3.6 V_H_ R98T and V_L_ L48Y mutations (38). The distinct State 2_V4_ HCDR3 orientations observed between the I5.6 and I3.6 suggested that these I3.6 mutations are effective due to germline-encoded HCDR3 residues. These HCDR3 residues, Y105, Y106, D107, and S109, are essential for germline-Env recognition (28) and, based on the kinetics data here, enhanced Env recognition in the I3.6 intermediate. The DMCs analysis revealed that these residues form highly networked long-range interactions that stabilize the bound state. The epitope protein peripheral residues Y106 and S109 showed ∼0.5-1.0 kcal/mol dissociation barrier couplings to Env K327, H330, and T415, likely mediated by structural stabilization of the HCDR3 loop at Y105 and backbone contacts with the epitope. These results underscore HCDR3 sequence specificity and suggest limited tolerance for HCDR3 residue substitution. This is consistent with substitutions at Y106 in a non-DH270.6 clade and at a terminal node splitting from I3.6, indicating weak neutralization breadth and potency compared to the mature DH270.6 bnAb. The DMCs also show that a non-bound-state salt bridge between Env GDIK motif K327 and HCDR3 D107 reduces the association barrier by 0.3 kcal/mol. This is consistent with observations in the MSM encounter ensemble, which show that this interaction stabilizes a near-bound state. These results, in conjunction with the MSM pathways, indicate that maturation in the DH270 clonal lineage tightly couples somatic mutations with germline-encoded motifs, focusing stabilizing contacts on conserved Env elements. More broadly, these findings suggest that immunogen design strategies targeting encounter-state geometry, rather than bound-state affinity alone, may accelerate the acquisition of breadth in otherwise difficult-to-induce antibody classes.

### Study Limitations

The structural states that make up the association pathway remain challenging to observe experimentally. Theoretical methods, therefore, remain the primary tool for investigating these states at high resolution. Our previous study demonstrated that MSMs are an effective immunogen design tool for selecting specific Env substitutions to confer enhanced association rates and, ultimately, improved selection of key breadth-conferring mutations in small-animal Ig-knock-in vaccination (38). These MSMs nevertheless have limitations. First, while sampling the forward association process is tractable, the reverse transition is often beyond the timescales accessible to current algorithms. This limits our ability to obtain a robust estimate of the equilibrium state distribution and, therefore, to provide a complete quantitative description of the interaction kinetics. Additionally, compressing the system’s large number of degrees of freedom into a few descriptive collective variables necessarily reduces state-to-state resolvability. Thus, we restrict the reported observables to relative locations, orientations, and average contact probabilities. These proved useful for immunogen design and were predictive of mutation patterns *in vivo* (38). DMCs are an established approach for evaluating the impact of paired residues on association transition barriers (50). In our previous study, a substitution that eliminated PNGS sites in the V4-loop at position 407 and at the epitope-proximal position 442 displayed strong coupling in the association barrier for I3.6, as predicted by the DH270.6 MSM. Similarly, here we show that HCDR3 D107_Ab_-K327_Env_ are coupled in the association barrier, as predicted by the I3.6 MSM. These provide a supportive predictive basis for the simulated ensemble. It is important to note, however, that the absence of coupling in the DMCs is not an indication that a residue pair does not show dependence. Standard errors ranged from 0.3-0.4 kcal/mol for measurements involving the dissociation barrier and 0.1-0.2 kcal/mol in the association barrier. Thus, residues whose contact coupling is weak or spread among multiple residues, as is often the case for flexible encounter states, cannot be reliably detected. Several experimental tools show promise for studying these transient states, including time-resolved SAXS (57), confocal single molecule Förster resonance energy transfer (58), and microsecond time-resolved cryo-electron microscopy (59). These advances, in combination with theoretical tools and mutagenesis, hold promise for shedding light on these important yet poorly understood structural states.

## Methods

### MD simulation

We prepared N- and C-terminal-truncated gp120 from the CH848 HIV-1 Env, isolated 949 days post-infection (PDB ID 6UM6) (60). The Fvs of I5.6 and I3.6 were prepared from the DH270.6 crystal structure (PDB ID 5TQA) (4, 61), reverting mutations using PyMOL (62). Each antibody was placed near the truncated CH848 gp120 using PyMol (62). The gp120 N-linked glycosylation sites were identified using the LANL glycosite server (63). The simulation systems were prepared using CHARMM-GUI Glycan Modeler (64) (65) (66) (67) (68). Each glycosylated complex was immersed in an octahedral water box with an edge length of 150 Å, and 0.15 M NaCl was added to neutralize the charge. The CHARMM36 force field and TIP3P water model (69) were used to run the MD simulation through pmemd CUDA in the AMBER22 MD simulation package (70) (71) (72). Hydrogen mass repartitioning (73) was used, allowing a larger simulation timestep. The total number of resulting atoms in each system was ∼75,000. The energy minimization was performed with 2,500 steps of steepest descent, followed by 2,500 steps of conjugate-gradient minimization. Equilibration and production runs were performed for 125 ps using a 1 fs timestep, followed by 250 ns production runs using a 4 fs timestep using the SHAKE algorithm (74) in the NVT ensemble. A total of 100 replicas of the MD simulation for each system.

### Adaptive MD simulation

The last frame of each production run was taken as the starting structure for adaptive molecular dynamics (MD) simulations. Adaptive MD was performed using the HTMD (High-Throughput Molecular Dynamics for Molecular Discovery) Python-based framework (75) in combination with the AMBER MD simulation package for the production simulations. In adaptive MD sampling, AMBER carried out classical MD production runs, while HTMD used the resulting trajectories to construct Markov state models (MSMs) to guide the selection of initial structures for subsequent simulation rounds. Trajectories were first projected onto a pairwise distance metric between interfacial residues. The selected residues and glycans are: gp120 - T135, V136, K137, N138, G139, T148, V149, P299, N300, N301, D321, I322, I323, G324, D325, I326, K327, Q327, A328, H330, N332, T415, Q417, antibody heavy chain - Y33, N52, T55, R57, N59, W101, G103, L104, Y105, Y106, D107, S108, S109, G110, Y111, N113, D115, antibody light chain - Y27, Y32, L48 (Y48 in I3.6 light chain), Y51, E52, P57, S58, Y93, S96, S97, and gp120 glycans - N138, N154, N301, N332, N408, N442. Based on this projection, a goal function defined as the inverse of RMSD to a reference bound structure (i.e., -RMSD) was applied to score each frame. The AdaptiveBandit algorithm (76) then selected frames with high goal scores (corresponding to low RMSD) as starting points for the next round of simulations, balancing exploitation of bound-like conformations with limited exploration of alternative states. These selected frames were then used as starting points for the next round of simulations. In total, 29 adaptive epochs were performed, each consisting of 100 replicas, with each replica running for 250 ns at NVT ensemble conditions.

### Markov State Model (MSM)

The final Markov State Model (MSM) was constructed using all the production MD trajectories generated from the adaptive sampling simulations. Trajectories were projected into a low-dimensional space based on the exponential pairwise distances between interfacial residues of the HIV-1 gp120 and the antibody, including glycan (I5.6: N138, N154, N332, N442, I3.6: N138, N154, N332, N408, N442) contacts observed in the bound complex. To capture the slowest collective motions associated with the binding process, time-lagged independent component analysis (TICA) (77) was applied with a lag time of 25 ns, retaining the top seven TICA dimensions. The resulting TICA space was then discretized using MiniBatch K-means clustering into 200 microstates. Finally, an MSM was constructed from the discretized trajectories using a lag time of 100 ns, based on the convergence of its plot, to identify the metastable states governing the binding dynamics.

### Ab-gp120 contact analysis

To understand the mechanism of N332 glycan capture followed by gp120 epitope recognition by I5.3 vs I3.6 HCRD3 loop, a contacts-based analyses were performed using all the MD simulation trajectories. For N332 glycan capture, contacts were calculated between the N332 glycan D-arm mannose residues and antibody V_H_ residues W101, Y111, N113, D115, Y116 and V_L_ residues L/Y48, Y51, E52, K55, R56, P57, S58. For gp120 epitope recognition by the HCRD3 loop, contacts between HCDR3 residues Y105, Y106, D107, S109, and gp120 glycan N332 base residues P299, H330, T415, Q417 and GDIK motif residues G324, D325, I326, K327 were analyzed. Contacts between selected residues were defined at a distance cutoff of 4 Å and computed using HTMD. The contacts were calculated for each macrostate of MSM, and the calculated contacts were weighted over all frames assigned to the macrostate of MSM. Different macrostates were quantified by the mean Cα atom RMSD, with the highest RMSD classified as the unbound state and the lowest RMSD as the bound state. In addition, variability in contacts within each macrostate was quantified by computing the standard deviation of the number of contacts per frame.

### Recombinant HIV-1 envelope SOSIP production

CH848 envelope SOSIPs were expressed in the Freestyle*TM* 293-F expression system (ThermoFisher Cat No. R79007). The Freestyle*TM* 293-F cells were diluted in Freestyle*TM* 293 Expression Medium (Cat No. 12338018) to 1.25x10*6* cells/mL. Plasmid DNA expressing each envelope SOSIP and furin were prepared in Kanamycin resistant LB broth cultures started from DH5α competent cell glycerol stocks and purified with Qiagen Plasmid Plus Kits (Cat No. at. No. 12981). The envelope SOSIP and furin were co-transfected at a 4:1 ratio and incubated with 293fectin*TM* transfection reagent (ThermoFisher Cat No. 12347019) in Opti-MEM I Reduced Serum Medium (ThermoFisher Cat No. 31985062) to allow for complex formation. The DNA-Fectin complex was added to the diluted 293F cell culture and incubated at 37°C, 8% CO*2* with orbital shaking at 120rpm for 6 days. Following incubation, the cell cultures were centrifuged at 4500 xg for 45 minutes and the cell supernatant harvested. The supernatant was filtered through a 0.45 µm PES filter and concentrated to approximately 100mL using a Vivaflow® 200 cross-flow cassette (Sartorius Cat No. VF20P2).

Purification of the expressed envelope SOSIP was performed using a PGT145 affinity chromatography column. The PGT151Ab affinity column was made by coupling PGT151 mAbs to CNBr-activated Sepharose 4B (Cat No. 170430-01, GE Bio-Sciences), packed into a Tricorn column (GE Healthcare) and equilibrated in 15mM HEPES, and 150mM NaCl (pH=7) buffer. The cell supernatant was run through the column at 2mL/min using an AKTA go chromatography system (Cytiva), followed by a buffer wash of two column volumes with 15mM HEPES, and 150mM NaCl. The envelope SOSIP was eluted off the column using 3M MgCl*2* and buffer exchanged into 15mM HEPES, and 150mM NaCl buffer. Buffer exchange was performed by ultrafiltration using 100 kDa MWCO Amicon® Ultra-15 Centrifugal Filter Units (Millipore Aldrich Cat No. UFC9010) and concentrated to <0.5 mL for size exclusion chromatography. Size exclusion chromatography was performed using a Superose 6 Increase 10/300GL Column (Cytiva) on an AKTA go system in 15 mM HEPES, and 150 mM NaCl. Fractions containing trimeric SOSIP were collected and subjected to quality control testing. Quality controls include: analytical SEC, SDS-PAGE, thermal shift analysis, biolayer interferometry (BLI), and negative stain electron microscopy (NSEM) to assure the presence of well-folded envelope trimers.

### Recombinant antibody production

Antibodies were expressed in the Expi293*TM* Expression System (ThermoFisher Cat No. A1435101). Expi293F*TM* cells were diluted to 2.5x10*6* cells/mL. Plasmid DNA for the antibody heavy and light chains were prepared in Kanamycin resistant LB broth cultures started from DH5α competent cell glycerol stocks and purified with Qiagen Plasmid Plus Kits (Cat No. at. No. 12981). Heavy and light chain plasmid DNA were co-transfected at a 1:1 ratio and incubated with Expifectamine 293 transfection reagent (ThermoFisher Cat. No. A14525) in Opti-MEM I Reduced Serum Medium (ThermoFisher Cat No. 31985062) to allow complex formation. The DNA-Expifectamine293 complex was then added to the prepared 293i cells and incubated at 37°C, 8% CO*2* with orbital shaking at 120rpm for 6 days. Following incubation, the cell cutures were centrifuged at 4500 xg for 45 minutes. The cell supernatant was harvested and filtered with 0.45 µm PES filter.

Purification of the expressed antibodies was performed using a Cytiva HiTrap Protein A HP purification column (Cat. No. 29048576). The column was equilibrated in 1X PBS buffer. The cell supernatant was run through the column at a flow rate of 0.5mL/min using an AKTA go chromatography system (Cytiva) followed by a buffer wash of two column volumes with 1X PBS. The antibody was eluted off the column with Pierce IgG Elution buffer, pH 2.8 (ThermoFisher Cat No. 21004). The sample elution was neutralized with Tris-pH 9 at 100µL/mL (ThermoFisher Cat. No. J60707). All antibodies produced were subjected to quality control testing including analytical SEC, SDS-PAGE and thermal shift analysis.

### Recombinant antibody Fab production

Antibody Fab were prepared using Lys-C digestion. IgG and Lys-C enzyme (ThermoFisher Cat. No. 90307) were combined in a 2000:1 ratio in 1X PBS. The digestion mixture was incubated at 37°C for 2 hours. Following the incubation, the sample was removed from the incubator, and a proteinase inhibitor was added to the solution to stop the digestion process. The antibody Fab are purified from the digestion mix with a Cytiva HiTrap Protein A HP purification column (Cytiva Cat No. 29048576). The full digestion mixture is applied to the column, and the Fab fragments are collected in the flow-through. The Fab are then buffer exchanged into 15mM HEPES and 150mM NaCl buffer. All Fab were subjected to quality control testing by SDS-PAGE and thermal shift analysis.

### Surface plasmon resonance

SPR binding measurements of DH270.6 and DH270 I3.6 Fabs against CH848 SOSIPs were performed using a Biacore T200 instrument (Cytiva) with HBS-EP+ 1X running buffer (Cytiva). PGT151 was immobilized on a CM5 sensor chip (Cytiva) to a response of 3500-5000 RU. CH848 SOSIPs were injected at 5 µL/min for a capture response of 200-500 RU. Fabs were injected over the captured SOSIP at 50 µL/min using the single cycle kinetics injection mode with a sequential series of 5 concentrations for 120s per concentration. The DH270.6, the unmutated I3.6, and Y106A & S109A mutant Fabs were measured at concentrations ranging 25-1350 nM with the Y105A & D107A mutant Fabs measured from 67.5-21600 nM. The final dissociation phase was conducted for 300-600s, with a 2-hour phase used to characterize the instrument detection threshold for interactions with immeasurable dissociation in 600s. The Fab and SOSIP were regenerated with three consecutive 10s pulses of 3M magnesium chloride at 100 µL/min. Fab binding to immobilized PGT151 as well as buffer binding to captured SOSIPs were used for double reference subtraction. Curve-fitting analysis was performed with the 1:1 binding model using the Biacore T200 Evaluation Software (Cytiva).

## Author Contributions

RCH conceptualized the study. SK performed MD simulations and Markov State modeling. CS prepared the protein for this study. YB performed Surface Plasmon Resonance. PA and JL assisted with SPR analysis. SK and RCH wrote the manuscript. All authors read and edited the manuscript.

## Data availability

All data generated in this study are available upon reasonable request.

## Acknowledgements

This study used the computational resources offered by Duke Research Computing (http://rc.duke.edu; NIH 1S10OD018164-01) at Duke University and was supported by NIH, NIAID, Division of AIDS Consortia for HIV/AIDS Vaccine Development (CHAVD) Grant UM1AI144371 (B.F.H.), R01AI145687 (P.A.), U54AI170752 (R.C.H. and P.A.), DP2-AI164323-03 (R.H.), and the Translating Duke Health Initiative (R.C.H. and P.A.).

