## Supplemental Information for "Germline-somatic residue synergy reshapes antibody encounter-state pathways to enhance HIV-1 recognition"

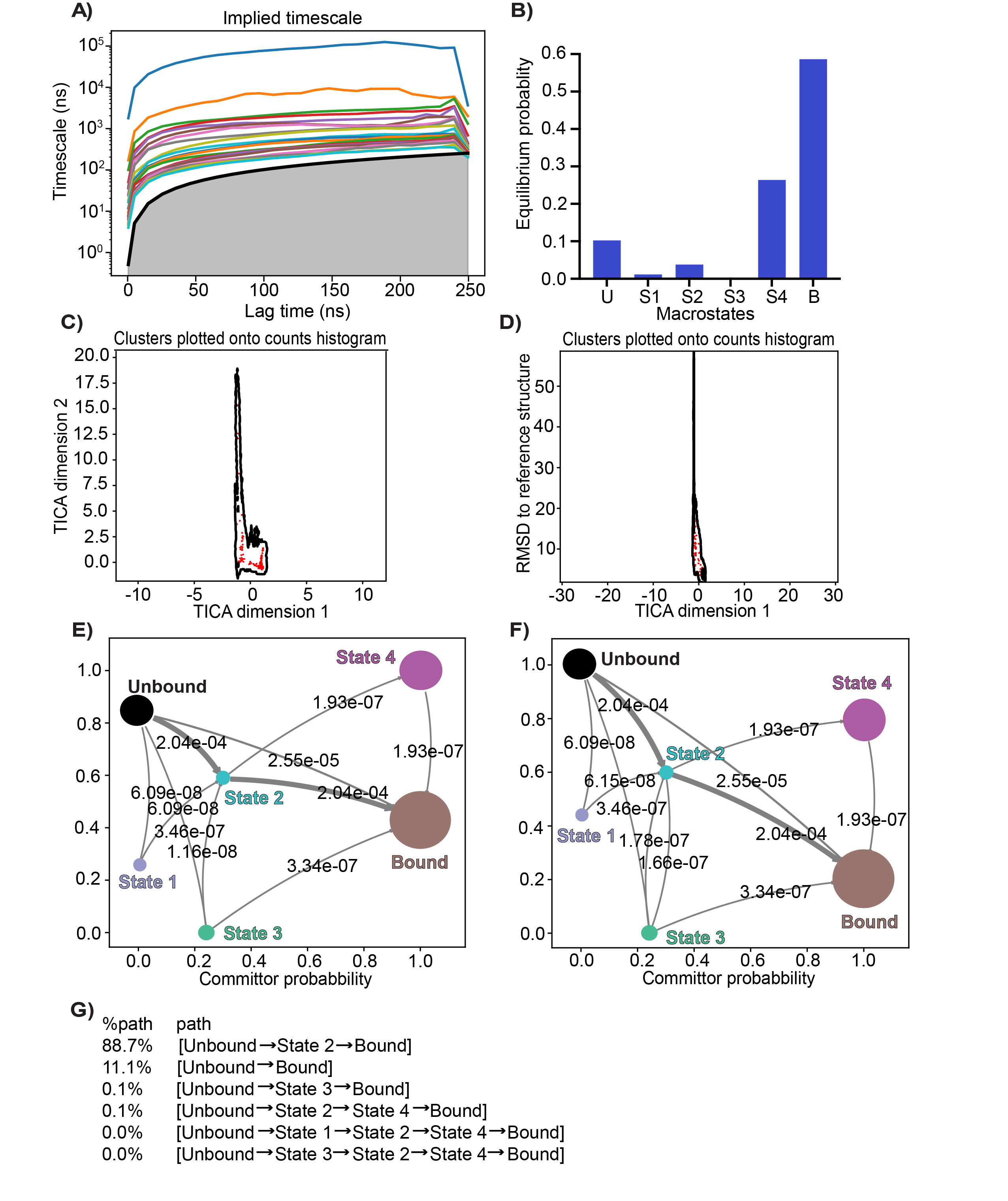
**Supplemental Figure 1.** Intermediate I5.6 association path modeling. **(A)** Clustered simulation set implied timescales plot. **(B)** MSM macrostate equilibrium probabilities. **(C)** Cluster centers overlayed on TICA dimension 1 and 2 distributions. **(D)** Cluster centers overlayed on TICA dimension 1 and RMSD distributions. **(E)** Committor probability plot depicting net flux from the unbound to bound macrostates. Circles are colored according to state, and circle sizes correspond to equilibrium probabilities. Arrows connect state and are sized according to transition flux with with overlayed flux values. **(F)** Gross flux between states depicted as in (*E*). **(G)** Path contributions to the association path.

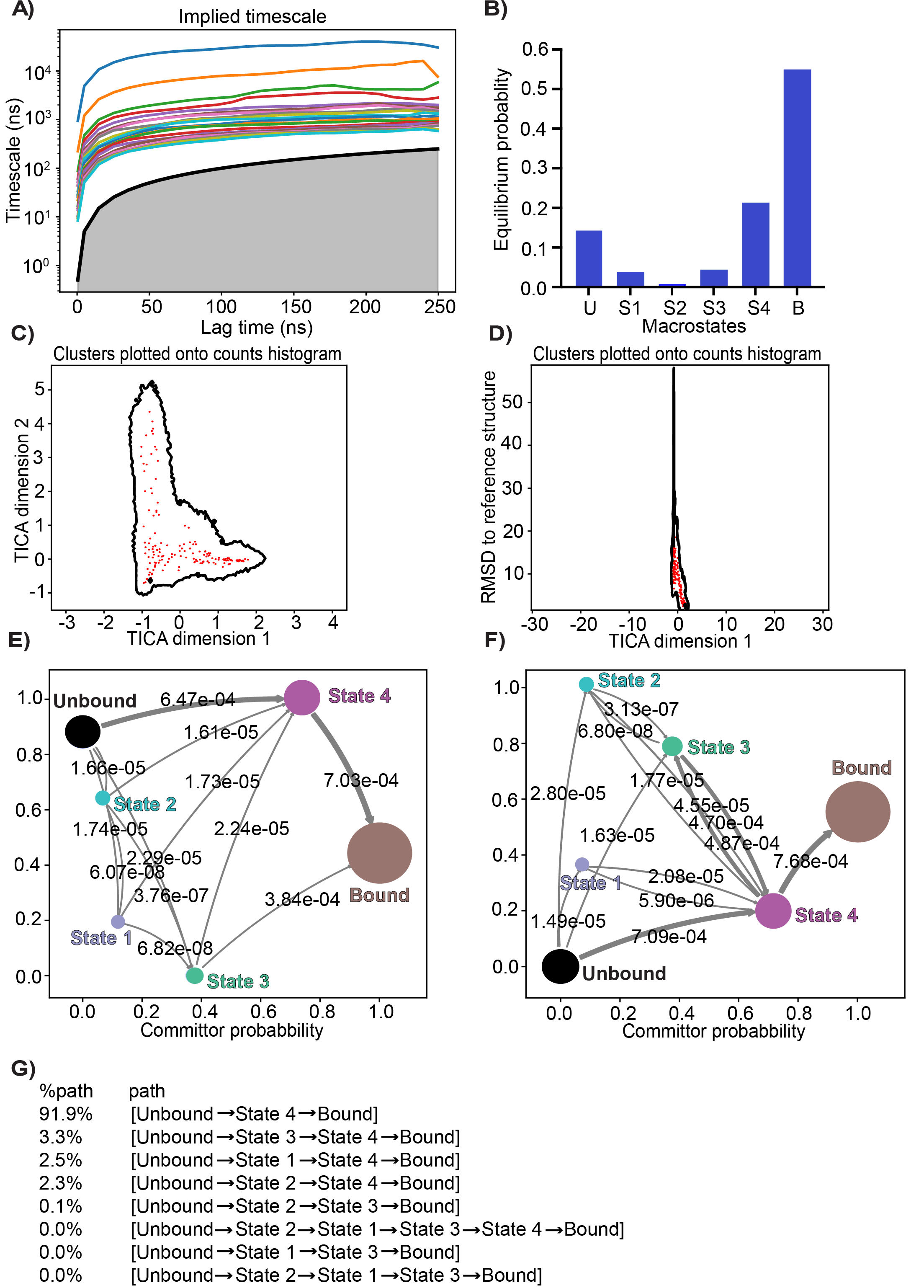
**Supplemental Figure 2.** Intermediate I3.6 association path modeling. **(A)** Clustered simulation set implied timescales plot. **(B)** MSM macrostate equilibrium probabilities. **(C)** Cluster centers overlayed on TICA dimension 1 and 2 distributions. **(D)** Cluster centers overlayed on TICA dimension 1 and RMSD distributions. **(E)** Committor probability plot depicting net flux from the unbound to bound macrostates. Circles are colored according to state and circle sizes correspond to equilibrium probabilities. Arrows connect state and are sized according to transition flux with with overlayed flux values. **(F)** Gross flux between states depicted as in (*E*). **(G)** Path contributions to the association path.

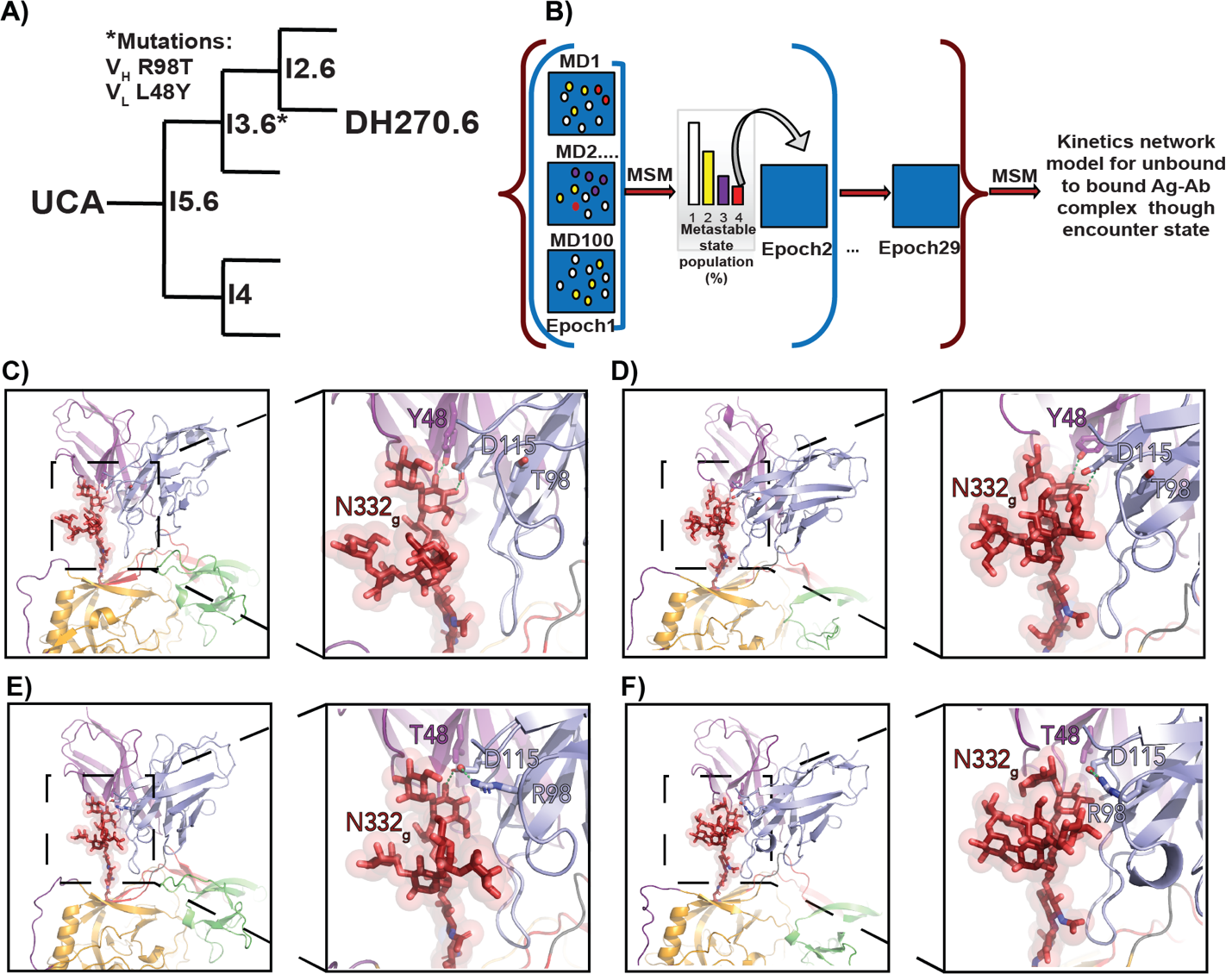
**Supplemental Figure 3. (A)** The DH270 clonal lineage beginning from the inferred germline (Unmuted Common Ancestor [UCA]), highlighting intermediates and the mature DH270.6 bnAb. The V_H_ R98T and V_L_ L48Y mutations that occur in I3.6 are indicated by the asterisk. **(B)** A schematic representation of Adaptive Sampling molecular dynamics simulations. **(C)** The experimentally determined I3.6 bound state structure (PDB ID 8SB1) highlighting interactions between the VH D115 and VL Y48 sidechain with the N332-glycan. **(D)** A representative bound state structure from the I3.6 molecular dynamics simulations. **(E)** The experimentally determined I5.6 bound state structure (PDB ID 8SAZ), highlighting interactions between the V_H_ D115 and V_L_ Y48 sidechain with the N332-glycan. **(F)** A representative bound state structure from the I5.6 molecular dynamics simulations.

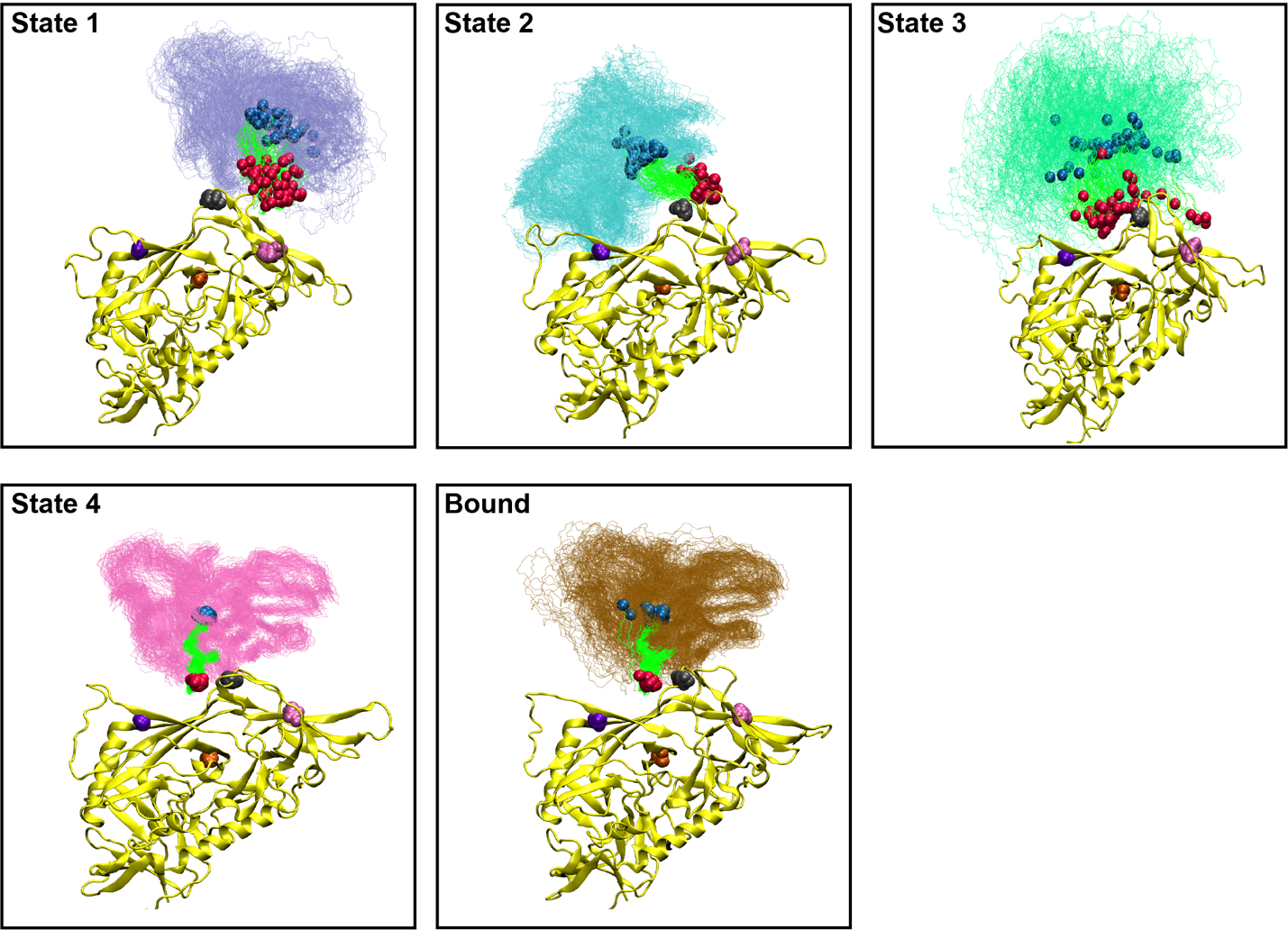
**Supplemental Figure 4.** I5.6 association path V_H_/V_L_ macrostate ensembles represented by 50 structures each (MSM weighted sampling) depicting the orientation of the HCDR3 loop (bright green). All gp120 domains were aligned such that ensembles represent distributions relative to gp120. A single gp120 structure is shown for visual clarity. The V4-loop (purple), gp120 core (orange), GDIK (grey), V1-loop base (pink), HCDR3 base (blue), and HCDR3 turn (red) centroid ensembles are represented as spheres.

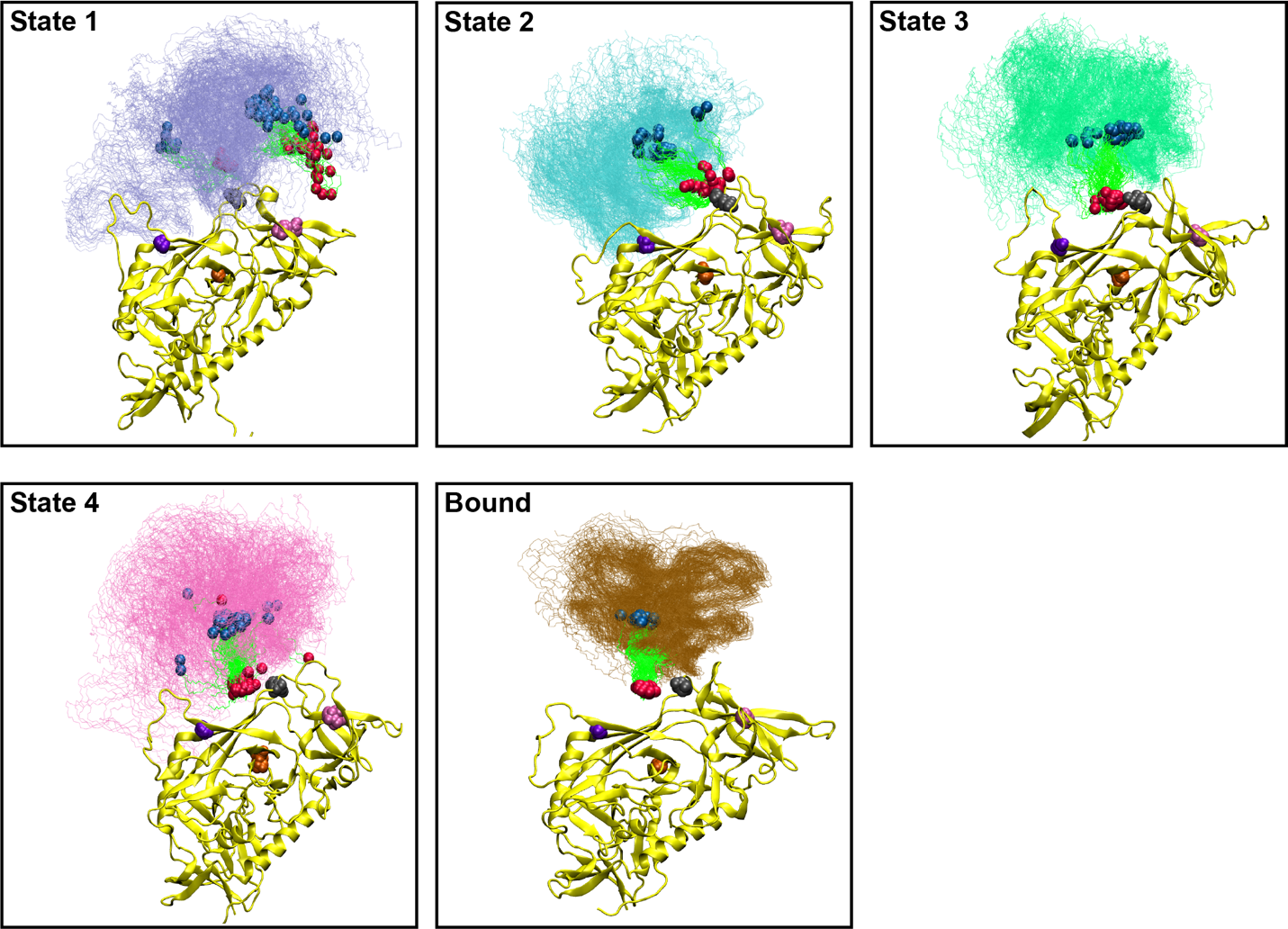
**Supplemental Figure 5.** I3.6 association path V_H_/V_L_ macrostate ensembles represented by 50 structures each (MSM weighted sampling) depicting the orientation of the HCDR3 loop (bright green). All gp120 domains were aligned such that ensembles represent distributions relative to gp120. A single gp120 structure is shown for visual clarity. The V4-loop (purple), gp120 core (orange), GDIK (grey), V1-loop base (pink), HCDR3 base (blue), and HCDR3 turn (red) centroid ensembles are represented as spheres.

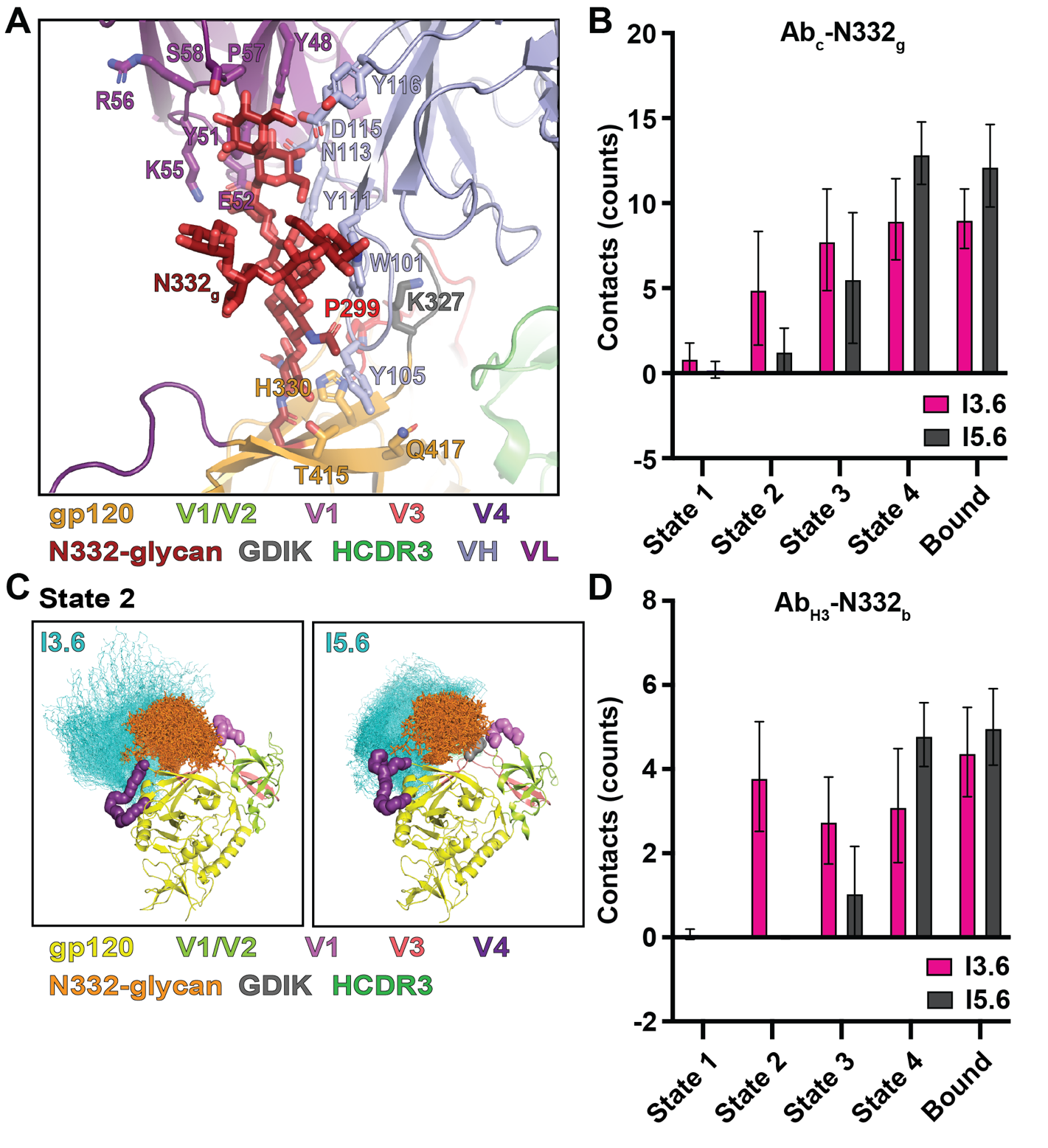
**Supplemental Figure 6. (A)** The bound state I3.6 structure (PDB ID 8SB1) depicting V_H_/V_L_ cleft and cleft proximal N332-glycan contact residues and HCDR3-epitope contact residues. **(B)** Weighted V_H_/V_L_ cleft and cleft proximal residue N332-glycan D-arm contact counts and standard deviations from the I5.6 and I3.6 MSMs. **(C)** State 2 V_H_/V_L_ macrostate ensembles represented by 50 structures each (MSM weighted sampling) depicting the N332-glycan ensemble distribution relative to the antibody ensemble. The gp120 domains were aligned as in *Supplemental Figure 4 and 5*. **(D)** Weighted HCDR3 residue epitope contact counts and standard deviations from the I5.6 and I3.6 MSMs.

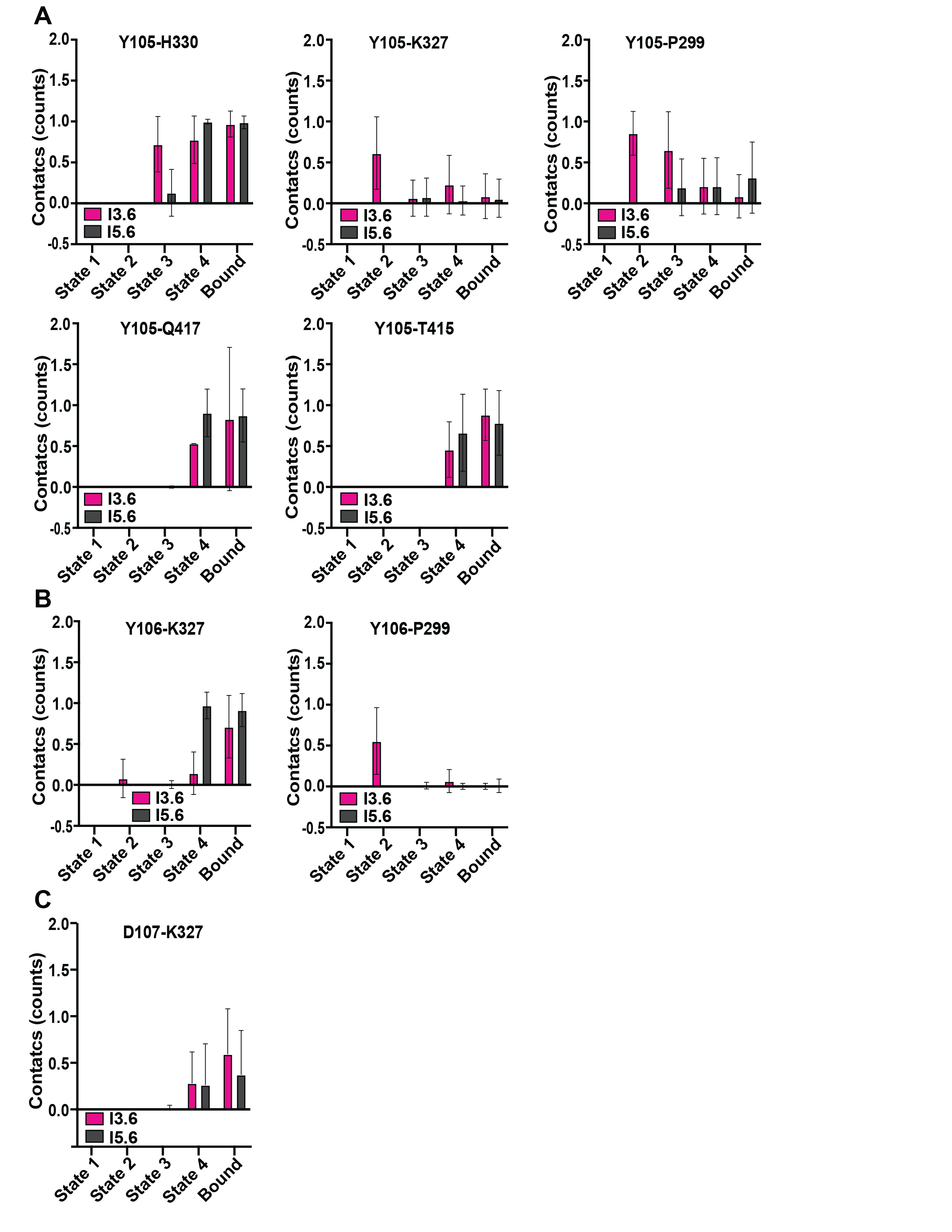
**Supplemental Figure 7. (A)** Weighted HCDR3 Y105 epitope contact counts and standard deviations from the I5.6 and I3.6 MSMs. **(B)** Weighted HCDR3 Y106 epitope contact counts and standard deviations from the I5.6 and I3.6 MSMs. **(C)** Weighted HCDR3 D107 epitope contact counts and standard deviations from the I5.6 and I3.6 MSMs.

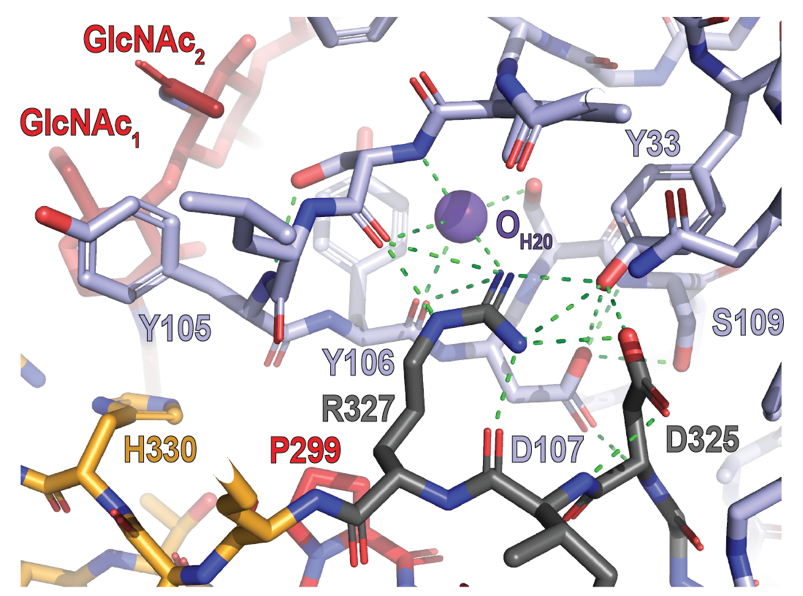

**Supplemental Figure 8.** High-resolution gp120 V3 peptide (orange/grey/red) bound DH270.6 structure (PDB ID 6CBP) depicting the contact network between GDIR residues (grey), HCDR3 (light blue), the N332-glycan (dark red) and a water molecule oxygen atom (purple).

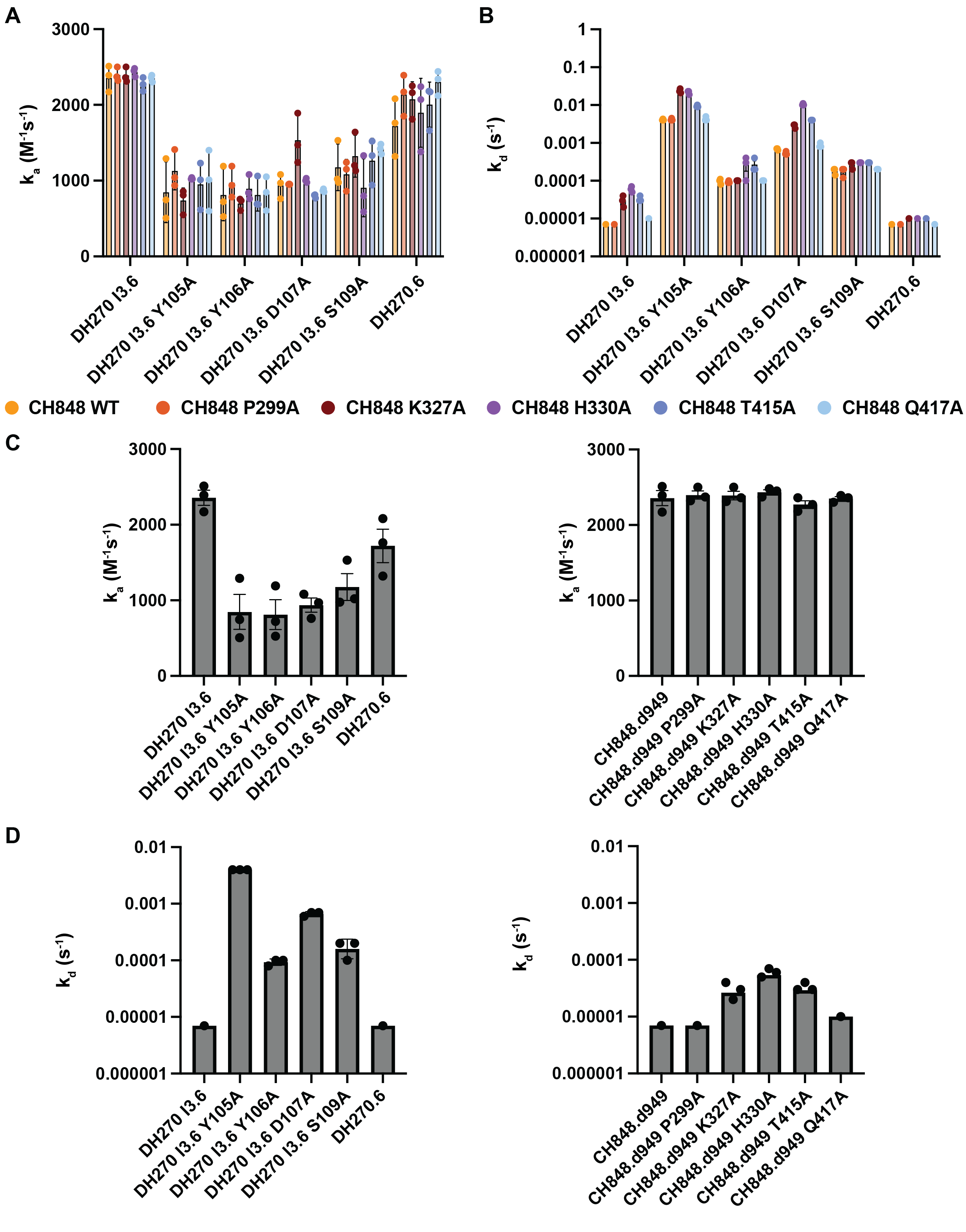
**Supplemental Figure 9. (A)** Mean association rate constants for all Fab and SOSIP constructs. Error bars indicate the standard error of the mean. All measurand values are shown (n=3). **(B)** Mean dissociation rate constants for all Fab and SOSIP constructs. **(C)** *(left*) Mean association rate constants for I3.6 Fab and I3.6 Fab mutants vs. the unmutated CH848.d949 SOSIP. (*right*) Mean association rate constants for I3.6 Fab vs. the unmutated CH848.d949 SOSIP and alanine-mutated CH848.d949 SOSIP constructs. **(D)** *(left*) Mean dissociation rate constants for I3.6 Fab and I3.6 Fab mutants vs. the unmutated CH848.d949 SOSIP. (*right*) Mean dissociation rate constants for I3.6 Fab vs. the unmutated CH848.d949 SOSIP and alanine-mutated CH848.d949 SOSIP constructs. (*A-D*) Error bars indicate the standard error of the mean. All measurand values are shown (n=1-4). Asterisks indicate values beyond the detection threshold (<7e-6 s^-1^).

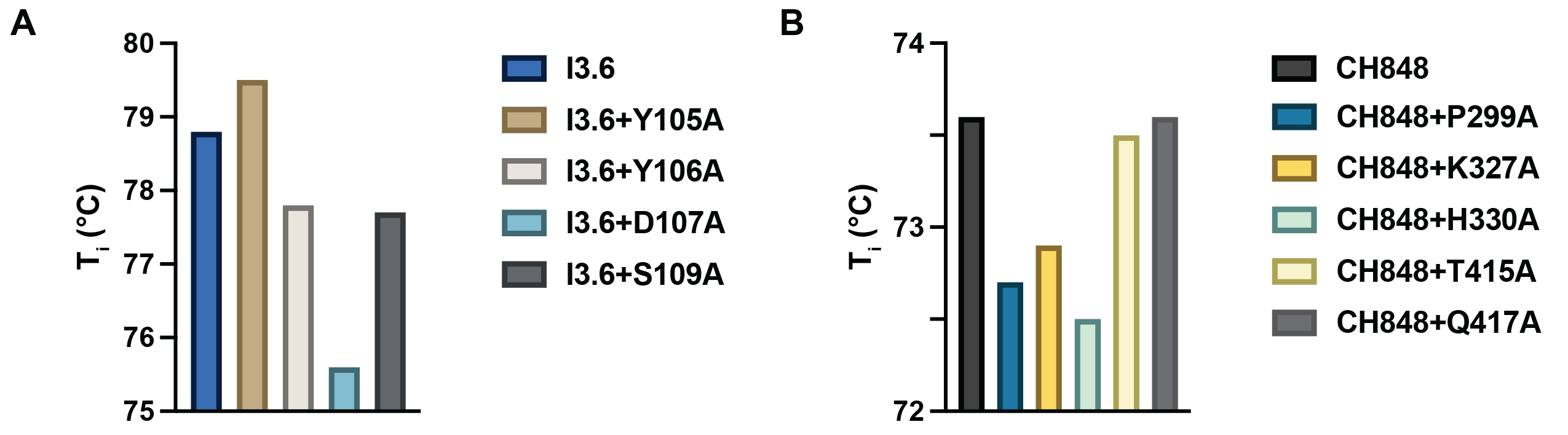
**Supplemental Figure 10. (A)** Thermal denaturation inflection temperatures (T_i_) for the I3.6 Fab and I3.6 alanine mutant Fabs. **(B)** Thermal denaturation inflection temperatures (T_i_) for the CH848 SOSIP and CH848 SOSIP alanine mutants.

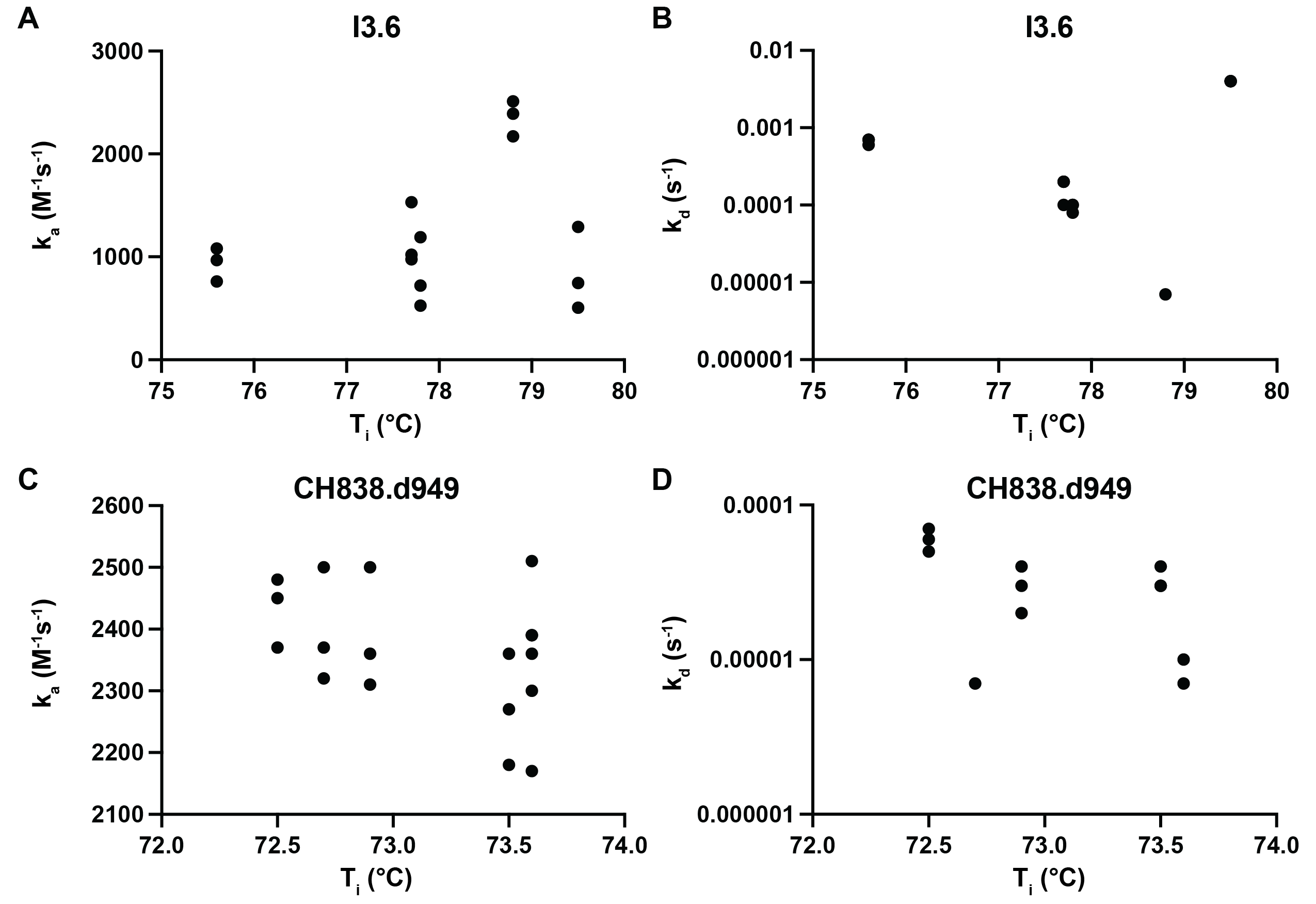
**Supplemental Figure 11. (A)** I3.6 and I3.6 alanine Fab mutant thermal denaturation inflection temperatures vs. CH848 SOSIP association rate constants, **(B)** I3.6 and I3.6 alanine Fab mutant thermal denaturation inflection temperatures vs. CH848 SOSIP dissociation rate constants, **(C)** CH848 SOSIP and CH848 SOSIP alanine mutant thermal denaturation inflection temperatures vs. I3.6 Fab association rate constants, and **(D)** CH848 SOSIP and CH848 SOSIP alanine mutant thermal denaturation inflection temperatures vs. I3.6 Fab dissociation rate constants. All measured values for each interaction pair are plotted in each panel (n=1-4).

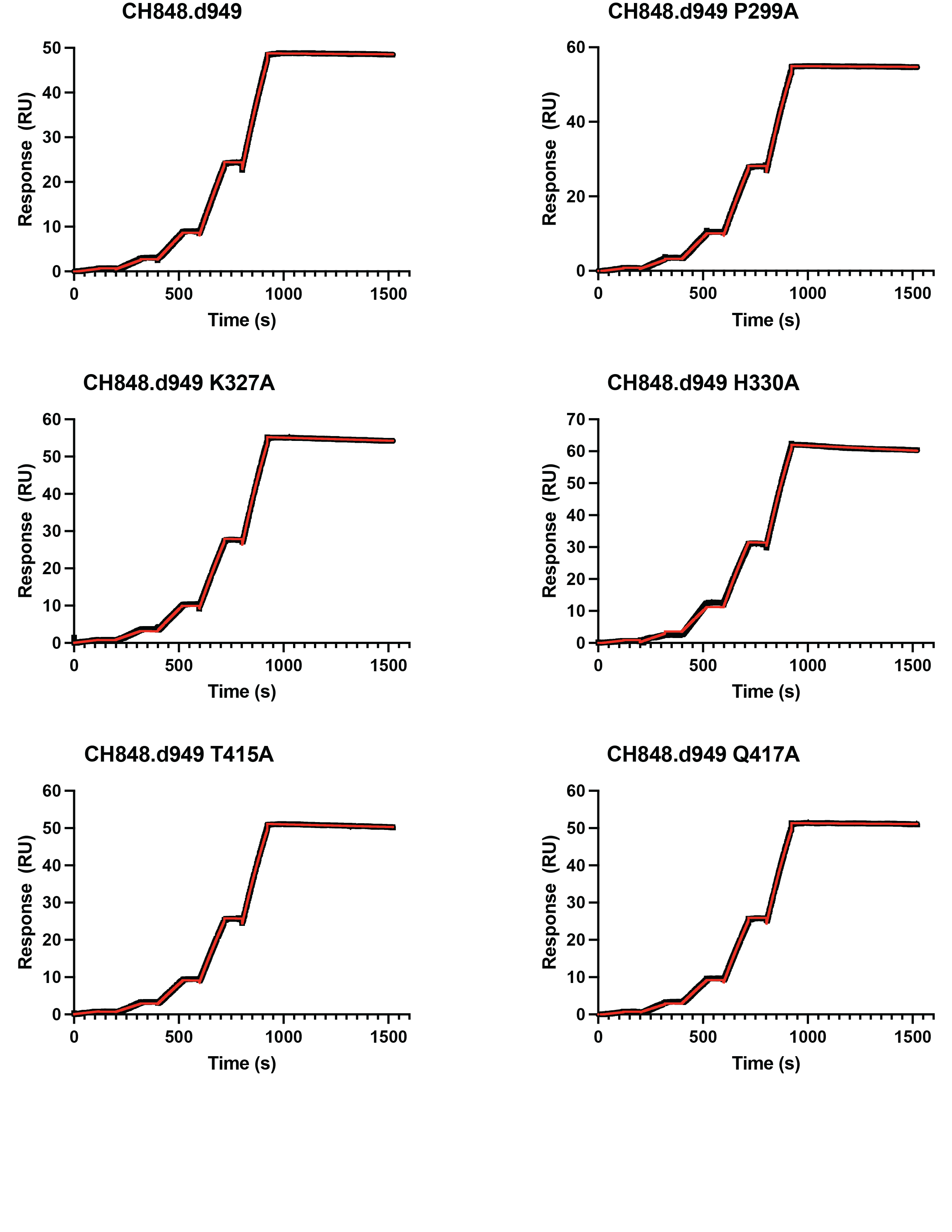
**Supplemental Figure 12.** Representative SPR curves (black) and fits (red) for the I3.6 Fab vs. CH848 SOSIP constructs.

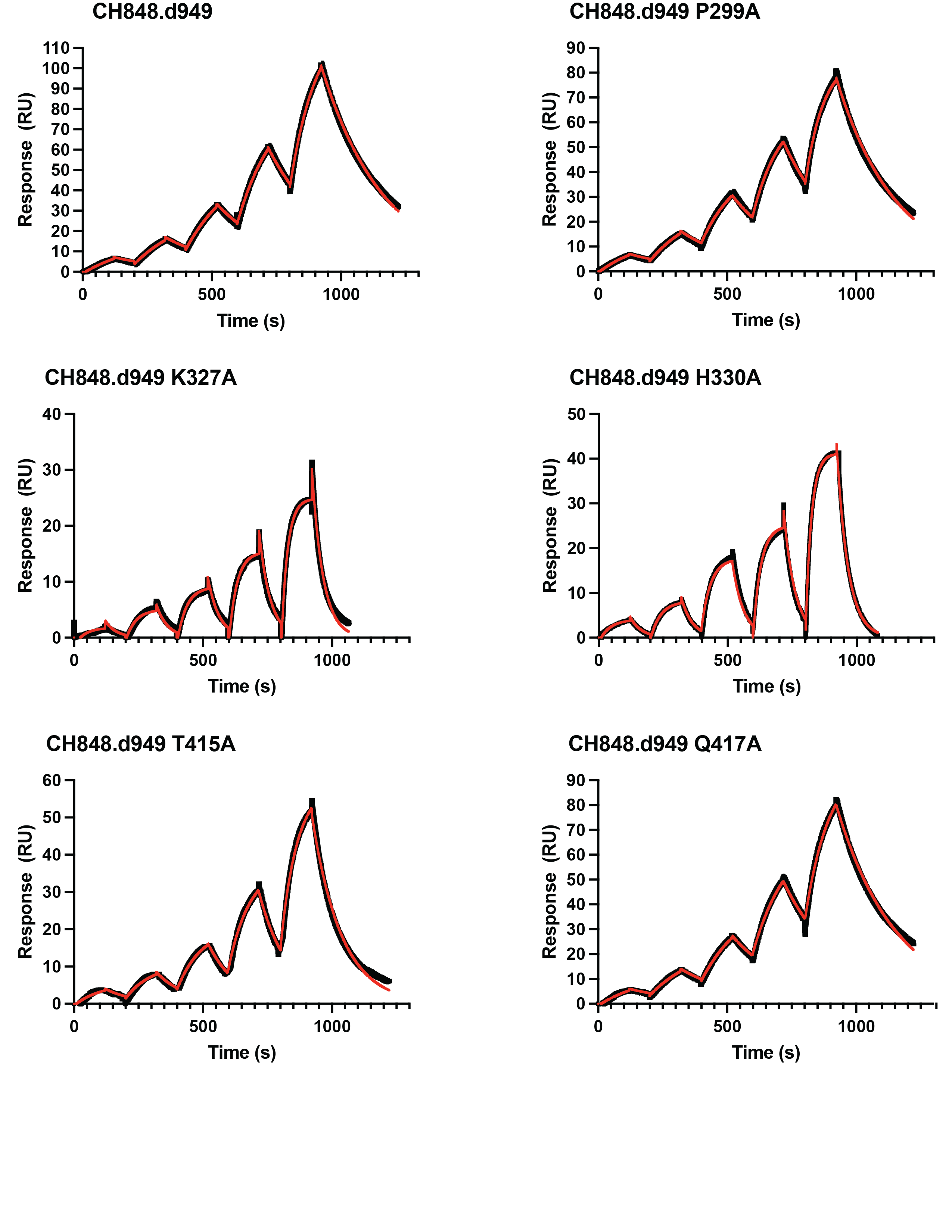
**Supplemental Figure 13.** Representative SPR curves (black) and fits (red) for the I3.6 Y105A Fab vs. CH848 SOSIP constructs.

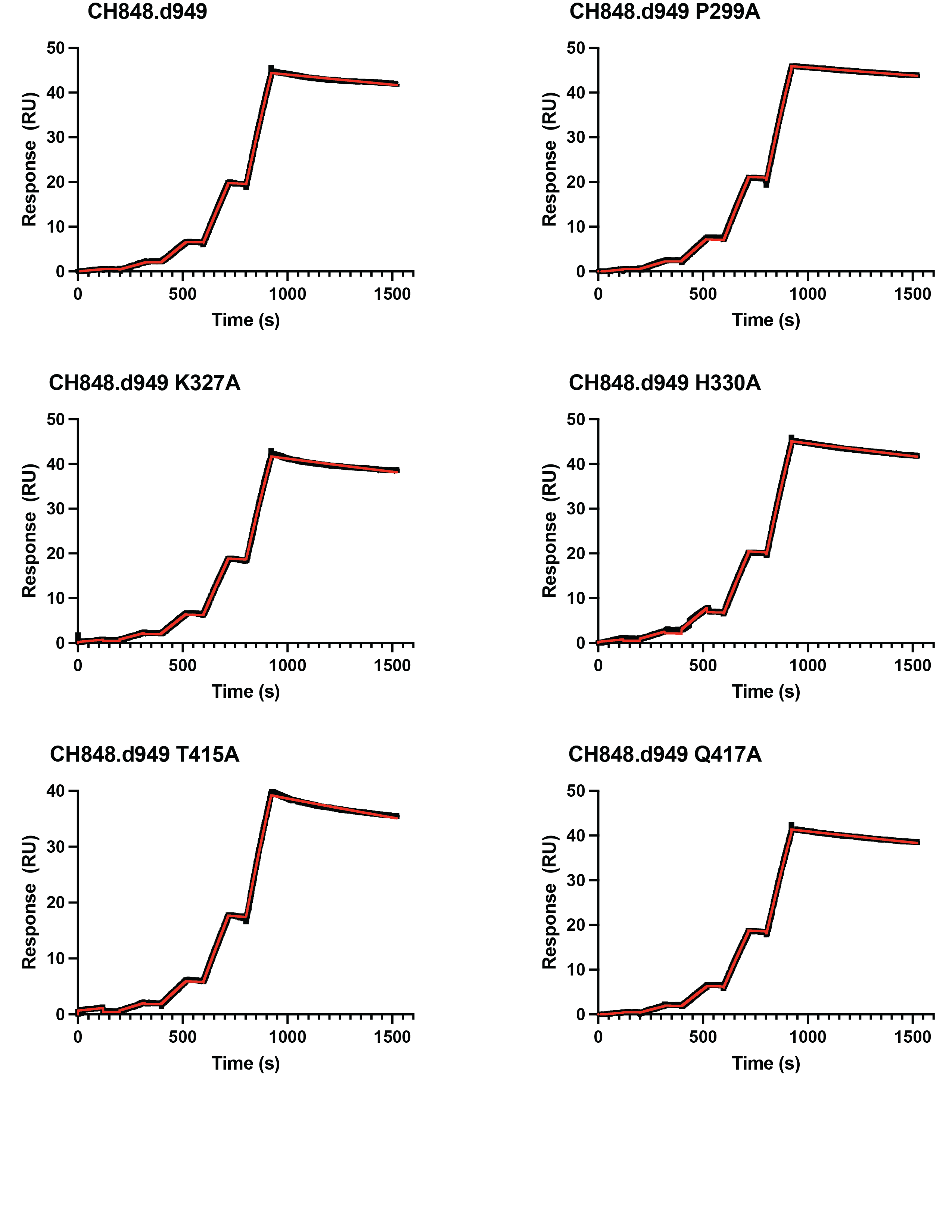
**Supplemental Figure 14.** Representative SPR curves (black) and fits (red) for the I3.6 Y106A Fab vs. CH848 SOSIP constructs.

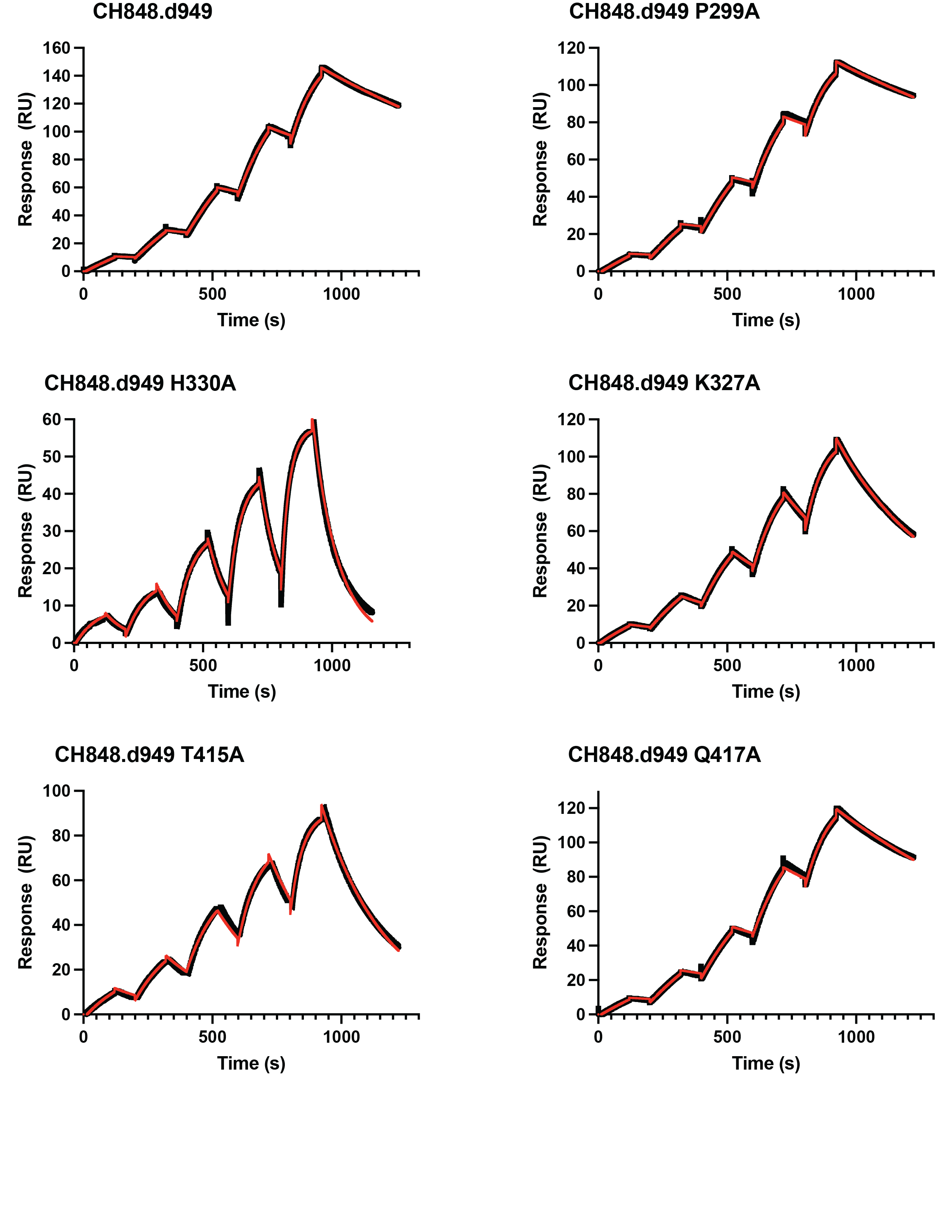
**Supplemental Figure 15.** Representative SPR curves (black) and fits (red) for the I3.6 D107A Fab vs. CH848 SOSIP constructs.

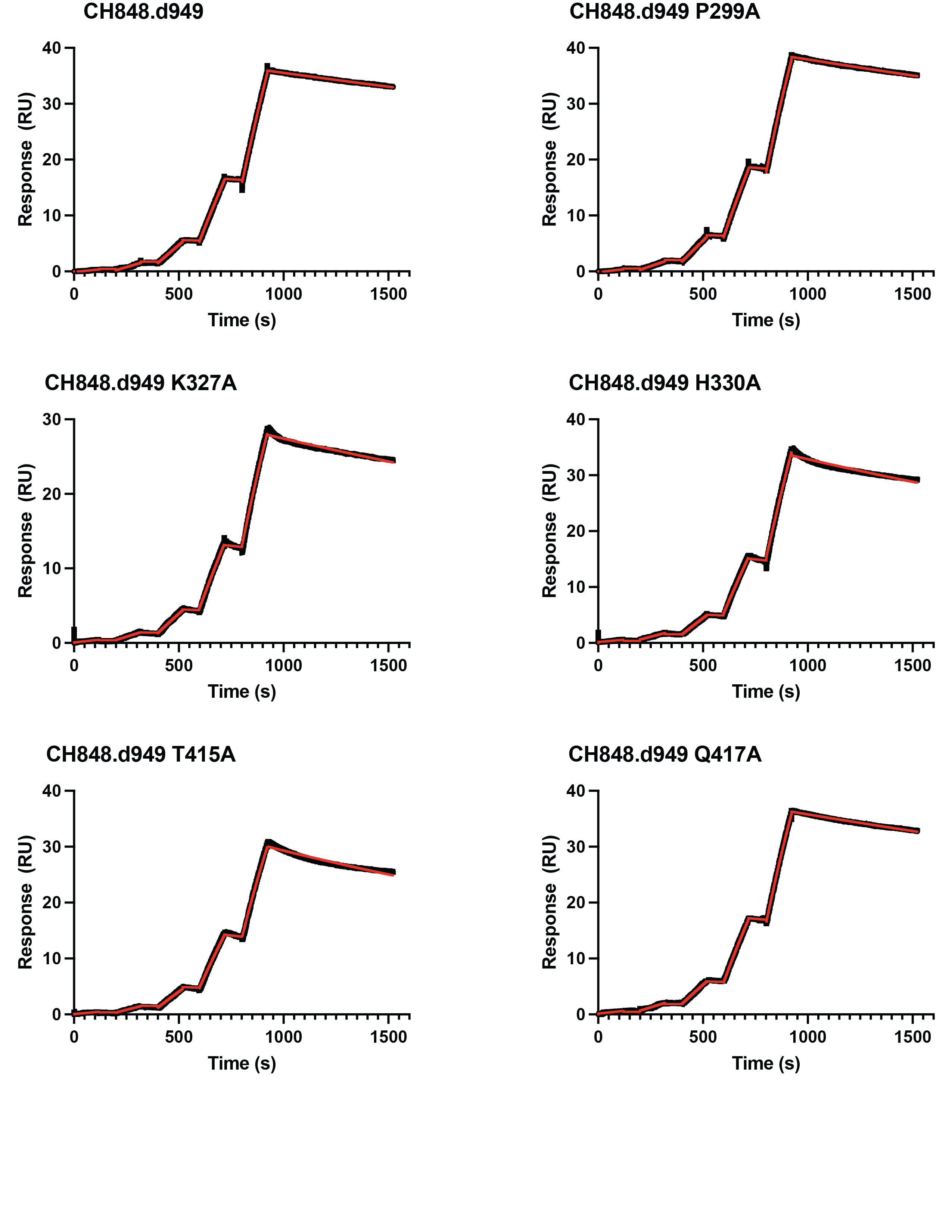
**Supplemental Figure 16.** Representative SPR curves (black) and fits (red) for the I3.6 S109A Fab vs. CH848 SOSIP constructs.

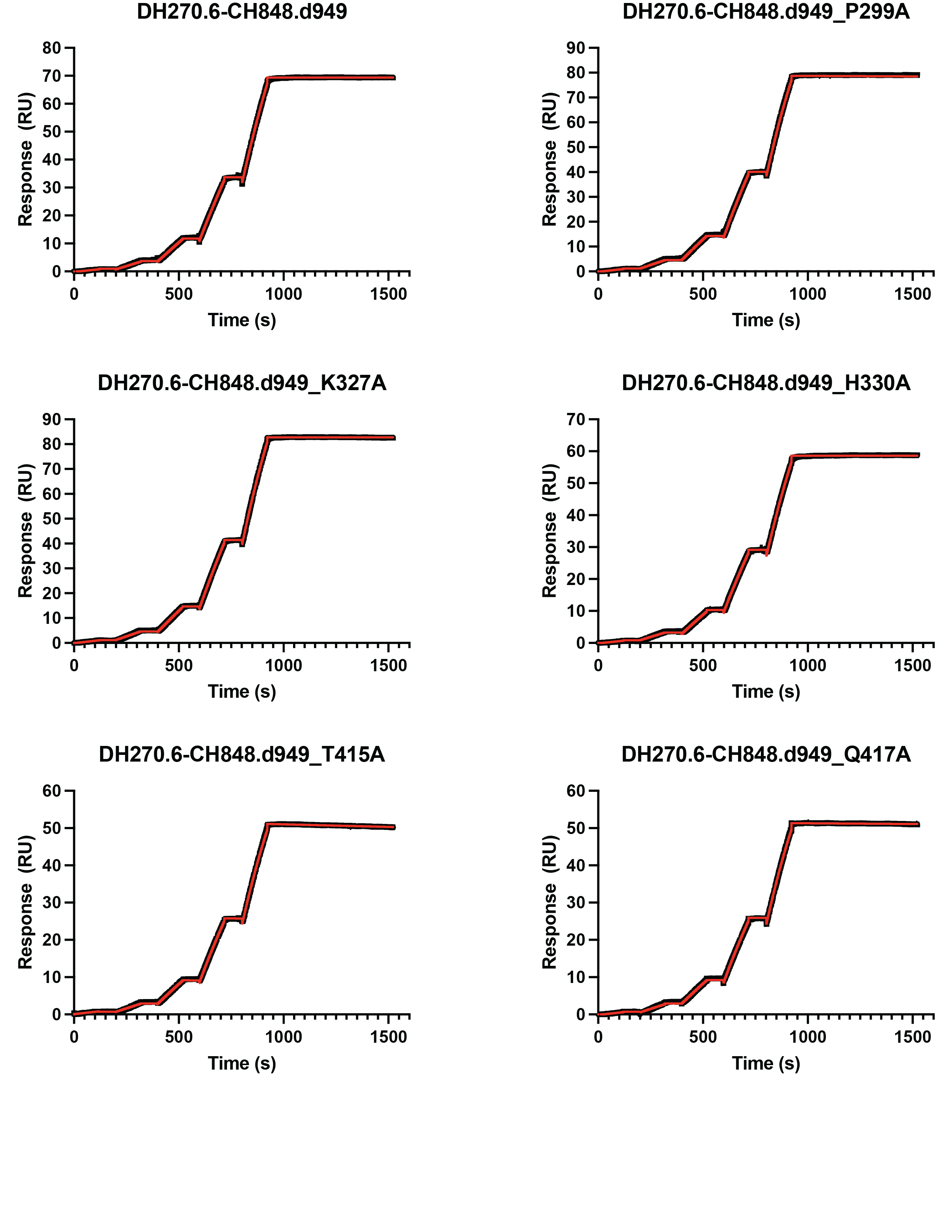
**Supplemental Figure 17.** Representative SPR curves (black) and fits (red) for the DH270.6 Fab vs. CH848 SOSIP constructs.
